# Proteome-wide neuropeptide identification using NeuroPeptide-HMMer (NP-HMMer)

**DOI:** 10.1101/2024.07.20.604414

**Authors:** Meet Zandawala, Muhammad Bilal Amir, Joel Shin, Won C. Yim, Luis Alfonso Yañez Guerra

**Affiliations:** Department of Biochemistry and Molecular Biology, University of Nevada, Reno, NV 89557, USA; Integrative Neuroscience Program, University of Nevada, Reno, NV 89557, USA; Neurobiology and Genetics, Theodor-Boveri-Institute, Biocenter, Julius-Maximilians-University of Würzburg, Am Hubland, 97074 Würzburg, Germany; School of Biology, University of Southampton, University Road SO17 1BJ, Southampton, UK; Institute for Life Sciences, University of Southampton, University Road SO17 1BJ, Southampton, UK

**Author notes:** Correspondence should be addressed to M.Z. and L.A.Y.G. Equal contribution (Meet Zandawala, Luis Alfonso Yañez Guerra).

**Keywords:** Neuropeptide, Hidden Markov model, Invertebrate, Evolution, Rotifer, Priapulid

## Abstract

Neuropeptides are essential neuronal signaling molecules that orchestrate animal behavior and physiology via actions within the nervous system and on peripheral tissues. Due to the small size of biologically active mature peptides, their identification on a proteome-wide scale poses a significant challenge using existing bioinformatics tools like BLAST. To address this, we have developed NeuroPeptide-HMMer (NP-HMMer), a hidden Markov model (HMM)-based tool to facilitate neuropeptide discovery, especially in underexplored invertebrates. NP-HMMer utilizes manually curated HMMs for 46 neuropeptide families, enabling rapid and accurate identification of neuropeptides. Validation of NP-HMMer on *Drosophila melanogaster, Daphnia pulex, Tribolium castaneum* and *Tenebrio molitor* demonstrated its effectiveness in identifying known neuropeptides across diverse arthropods. Additionally, we showcase the utility of NP-HMMer by discovering novel neuropeptides in Priapulida and Rotifera, identifying 22 and 19 new peptides, respectively. This tool represents a significant advancement in neuropeptide research, offering a robust method for annotating neuropeptides across diverse proteomes and providing insights into the evolutionary conservation of neuropeptide signaling pathways.

## Introduction

Neuropeptides and peptide hormones are the largest class of neurochemical messengers. They regulate diverse animal behaviors and physiological processes including feeding, locomotion, metabolism, growth, and reproduction (Nässel and Zandawala, 2019). Consequently, they are implicated in various diseases ranging from obesity and diabetes to high blood pressure and sleep disorders. Further, the origins of several neuropeptide families can be traced back to the common ancestor of Bilateria (Jekely, 2013; Mirabeau and Joly, 2013). Thus, orthologs of several vertebrate neuropeptide families are found in invertebrates such as insects, nematode and polychaete worms, and echinoderms. These include orthologs of vertebrate neuropeptides like calcitonin, galanin, orexin, neuropeptide-S, gonadotropin-releasing hormone (GnRH), thyrotropin-releasing hormone (TRH), corticotropin-releasing factor, glycoprotein hormones, insulin, substance P, neuropeptide Y, and vasopressin/oxytocin, amongst others (Bauknecht and Jekely, 2015; Istiban et al., 2024; Lindemans et al., 2009; Nässel and Zandawala, 2019; Nässel et al., 2019; Odekunle et al., 2019; Semmens et al., 2015; Stafflinger et al., 2008; Staubli et al., 2002; Tian et al., 2016; Van Sinay et al., 2017; Wegener and Chen, 2022; Zandawala, 2012). In many instances, the functions of these signaling systems have also been conserved throughout evolution (Nässel and Zandawala, 2019; Nässel et al., 2019; Van Sinay et al., 2017). Therefore, discovering and characterizing neuropeptide signaling pathways in invertebrates can offer valuable insights into their vertebrate orthologs.

Despite their widespread occurrence and medical significance, establishing evolutionary relationships between neuropeptides across different animal phyla has historically been quite challenging due to the short sequences of biologically active mature peptides. Additionally, bioinformatics approaches such as Basic Local Alignment Search Tool (BLAST) are often unsuitable for discovering neuropeptide precursors in animals belonging to phyla other than Arthropoda, Nematoda, Annelida, Mollusca, and Chordata, where many neuropeptides have already been identified. This limitation is particularly pronounced for neuropeptide precursors that generate mature peptides with limited conservation beyond the phyla in which they were originally discovered. For instance, although the structure of vertebrate TRH (comprised of only 3 amino acids) was discovered in 1969 (Boler et al., 1969; Burgus et al., 1969), it was not until 2015 that the identity of invertebrate TRH was revealed through the functional characterization of *Platynereis dumerilii* TRH receptors (Bauknecht and Jekely, 2015). While this issue can be circumvented by using neuropeptide prediction tools based on machine learning models (Agrawal et al., 2019; Kang et al., 2019; Ofer and Linial, 2014; Wang et al., 2023; Wang et al., 2024), these existing tools are unable to determine the neuropeptide family to which the predicted neuropeptide precursor belongs. The large number of neuropeptide families, some of which have similar conserved motifs, further complicates accurate classification of neuropeptides.

To address this gap, we have developed NeuroPeptide-HMMer (NP-HMMer), a hidden Markov model (HMM)-based tool designed to facilitate neuropeptide discovery, particularly in understudied arthropods and other invertebrate phyla. NP-HMMer is based on manually curated HMMs for 46 invertebrate neuropeptides, which enables rapid and accurate identification of neuropeptides in entire proteomes by presenting output in multiple forms (i.e., trimmed and non-trimmed sequence alignments, and sequence logos). We demonstrate the effectiveness of this tool by discovering members of several neuropeptide families in Priapulida (penis worms) and Rotifera (wheel animalcules).

## Methods

### Identification and curation of neuropeptide precursors for HMM generation

Members of 46 neuropeptide precursor families were used as queries to search for orthologs from proteomes of the following 17 representative arthropod species: *Apis mellifera, Aedes aegypti, Bombyx mori, Acyrthosiphon pisum, Daphnia pulex, Drosophila melanogaster, Ixodes scapularis, Lepeophtheirus salmonis, Nasonia vitripennis, Locusta migratoria, Pediculus humanus, Rhodnius prolixus, Strigamia maritima, Tetranychus urticae, Tribolium castaneum, Varroa destructor, and Zootermopsis nevadensis*. These species encompass a broad phylogenetic range that includes insects, crustaceans, myriapods, and chelicerates. Neuropeptide precursors were identified using BLASTp with an E-value threshold of 1e-2 to ensure the identification of orthologs. The search was limited to the top three hits for each query sequence and the top high-scoring pair for each target sequence. Following BLASTp, the sequences were manually curated to remove redundant or partial sequences. Sequences not encoding neuropeptides were also removed by manually scanning for characteristic prohormone cleavage sites and conserved motifs in mature peptides. Signal peptides were identified using SignalP 6.0 (Teufel et al., 2022).

### NP-HMMer generation and validation

Ortholog sequences belonging to the same neuropeptide family were aligned using MUSCLE (Edgar, 2004), followed by manual refinement to optimize the positioning of conserved residues and motifs. HMMER software suite (http://hmmer.org/) was then used to construct and train family-specific HMMs using the refined sequence alignments. HMMs were trained using default parameters and the hmmbuild command, with a minimum of five neuropeptide precursor sequences used for each model. Proteomes of *Drosophila melanogaster, Daphnia pulex* and *Tribolium castaneum* were scanned using the trained HMM profiles with hmmsearch and an E-value of 1e-2. Identified sequences were manually inspected to confirm characteristic neuropeptide features for each family. In most of the cases, the models identified the correct neuropeptide. In cases where the model did not obtain accurate hits, sequence alignments were modified, and models were generated.

### Neuropeptide identification in Priapulida and Rotifera

Validated models were used to identify neuropeptide precursors in whole predicted proteomes of one priapulid (*Priapulus caudatus*, NCBI; GCF_000485595.1) and four rotifers ((*Adineta ricciae* (UNIPROT; UP000663828), *Brachionus plicatilis* (UNIPROT UP000276133), *Didymodactylos carnosus* (UNIPROT; UP000663829) and *Rotaria socialis* (UP000663873)) using an E-value of 1e-2, and -domE e1-6.

### Sequence annotation and visualization

Signal peptides were predicted using SignalP 6.0. Monobasic and dibasic cleavage sites, and mature peptides were manually annotated based on the NP-HMMer output. Sequence alignments in the manuscript were generated using Clustal Omega (https://www.ebi.ac.uk/jdispatcher/msa/clustalo) with default settings. The alignments were shaded using Boxshade (https://junli.netlify.app/apps/boxshade/) based on at least 60% amino acid conservation as the minimum for highlighting.

## Results and discussion

### NP-HMMer generation

HMM-based searches can be powerful in discovering orthologs of genes that evade BLAST-based searches. This is due to their ability to model positional dependencies between amino acids, allowing HMMs to capture conserved motifs interrupted by insertions or deletions. HMMs use a probabilistic framework to represent amino acid distributions, accommodating variability within a family while still detecting distant homologs. Furthermore, HMMs can be used to construct profile models that represent the consensus of a protein family, enabling the identification of new family members with limited sequence similarity to known members. We previously used this approach to identify GnRH in the amphioxus *Branchiostoma floridae* (Yanez Guerra and Zandawala, 2023). In contrast, BLAST, relying on pairwise alignments, may struggle to detect relationships when sequence similarity is low (Eddy, 2004). Hence, we have generated NP-HMMer **(Figure 1)**, a tool that allows annotation of 46 invertebrate neuropeptides in whole proteomes. NP-HMMer is based on python and the open-source version is freely-available with step-by-step instructions for setup and execution locally.

**Figure 1:**
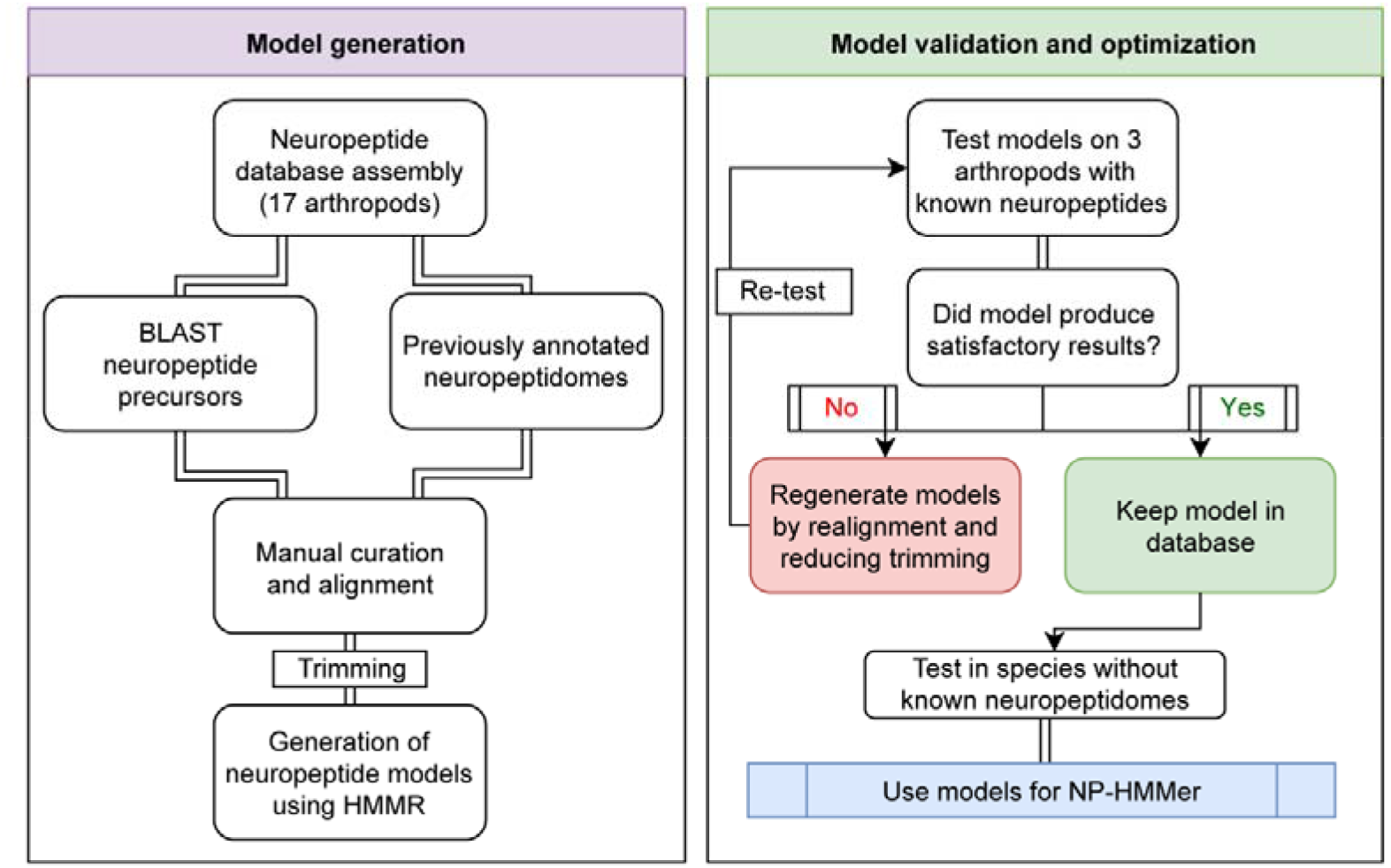
Workflow for creating the NP-HMMer search tool. A database of neuropeptide precursors from up to 17 representative arthropod species was first assembled. This was achieved using a combination of BLAST analyses and by manually searching previously annotated neuropeptidomes. Complete precursor sequences belonging to specific neuropeptide families were curated and aligned. The resulting alignments were trimmed to exclude non-conserved regions and HMMs were generated. The models were tested using 3 arthropod proteomes whose neuropeptidomes were previously established. Models producing accurate predictions were kept, while others were refined and retested. The final models formed the basis of NP-HMMer which was used to identify neuropeptides in priapulids and rotifers.

### NP-HMMer validation

We validated our models by testing their effectiveness in identifying neuropeptides in proteomes of *Drosophila melanogaster* **(Figure S1)**, *Daphnia plex* **(Figure S2)** and *Tribolium castaneum* **(Figure S3)**, whose neuropeptidomes have been previously established (Dircksen et al., 2011; Hewes and Taghert, 2001; Li et al., 2008; Nässel and Zandawala, 2019). Our search identified members of all neuropeptide families in these species **(Table S1)**. *Daphnia* diuretic hormone 44 (DH44) and myosuppressin, as well as *Tribolium* natalisin were missing in the proteomes used for the analysis. Hence, manual addition of these neuropeptide sequences to their respective proteomes enabled their detection by NP-HMMer. Since sequences from *Drosophila, Daphnia* and *Tribolium* were used to generate the models, we also tested the effectiveness of NP-HMMer using a proteome from *Tenebrio mollitor* whose sequences were not used to generate NP-HMMer. We were able to retrieve all the neuropeptides previously identified in *Tenebrio* (Veenstra, 2019) for which the models are available **(Table S2)**. In addition, we also identified additional neuropeptide isoforms and transcript variants not identified previously. Hence, NP-HMMEr is effective at identifying neuropeptides in a wide-range of insects.

In some cases, our models could not distinguish between closely-related neuropeptides. For example, allatostatin-C, allatostatin-CC, and allatostatin-CCC have similar signatures that prevent automatic classification (Veenstra, 2016). Similarly, adipokinetic hormone (AKH), corazonin and AKH/corazonin-related peptide (ACP), bursicon alpha and bursicon beta, CCHamide-1 and CCHamide-2, as well as natalisin and tachykinin are quite similar which prevents unambiguous identification. However, manual examination of the search results, which are conveniently presented in the form of sequence alignments and logos **(Figure S4)**, can readily resolve any discrepancies. We also provide a reference guide to facilitate the identification of these closely-related neuropeptides **(Figure S5)**. Regardless, this approach is still effective as we were able to identify phoenixin precursor **(Figure S1)** for the first time in *Drosophila melanogaster*.

### Neuropeptide identification in Priapulida and Rotifera

Having validated NP-HMMer, we decided to comprehensively identify neuropeptide complements of Priapulida and Rotifera. Previous studies have independently identified orthologs of AKH-like (Li et al., 2016), neuropeptide F (Yanez-Guerra et al., 2020), RYamide/Luqin (Yanez-Guerra et al., 2018), pigment-dispersing factor (Mayer et al., 2015), ion transport peptide (Gera et al., 2024), vasopressin/oxytocin (VP/OXT) (Lockard et al., 2017), allatostatin-CC (misclassified as allatostatin-C) and FMRFamide-like peptides (Christie et al., 2011) in priapulids. In contrast, to the best of our knowledge, only AKH-like precursor has been discovered in rotifers (Cadena-Caballero et al., 2023; Hauser and Grimmelikhuijzen, 2014). Hence, we chose species from these two phyla to assess the suitability of NP-HMMer in discovering orthologs of arthropod neuropeptides in other phyla. Specifically, we mined proteomes of one priapulid (*Priapulus caudatus* **(Figure S6)**) and four rotifers (*Adineta ricciae* **(Figure S7)**, *Brachionus plicatilis* **(Figure S8)**, *Didymodactylos carnosus* **(Figure S9)** and *Rotaria socialis* **(Figure S10)**) using NP-HMMer. Our search identified members of 25 and 20 neuropeptide families in Priapulida and Rotifera, respectively (**Table 1, Figures 2-3 and S11**). When compared to previously discovered neuropeptides in these phyla, this represents 22 and 19 novel neuropeptides in Priapulida and Rotifera, respectively. In particular, we identified orthologs of crustacean cardioactive peptide (CCAP) **(Figure 2B)**, leucokinin **(Figure 2C)**, allatostatin-A **(Figure 2F)**, proctolin **(Figure 2H)**, allatostatin-CCC **(Figure 2I)**, pigment-dispersing factor (PDF) **(Figure 2L)**, VP/OXT **(Figure 2N)** FMRFamide **(Figure 3C)**, CCHamide **(Figure 3D)**, CAPA **(Figure 3E)**, neuropeptide F **(Figure 3H)**, orcokinin **(Figure 3I)**, phoenixin **(Figure 3J)**, bursicon alpha, glycoprotein hormone alpha 2 (GPA2) and insulin-like peptides **(Figure S11)** for the first time in rotifers. We also identified precursors encoding two copies of Wamides in *Adineta* **(Figure S7)** and *Didymodactylos* **(Figure S9)**. These peptides appear to be the ancestral form of allatostatin-B as loss of a cleavage site in between the two *Adineta* Wamides could give rise to an allatostatin-B-like peptide **(Figure 3F)**. Moreover, we identified a novel AKH-like precursor in *Brachionus*. Sequence alignment suggests that this AKH is more similar to other arthropod AKH peptides, whereas the AKH-like peptide discovered previously (Hauser and Grimmelikhuijzen, 2014) appears to be more similar to arthropod ACP **(Figure 3G)**. Functional characterization of AKH and ACP receptors in the future is needed to unambiguously determine their ligands. This limitation also holds true for all other neuropeptides identified here as they can only be considered predictions until functionally characterized. Lastly, we also identified a calcitonin ortholog in *Priapulus* which is significantly shorter compared to other arthropod calcitonin **(Figure 3A)**. Additional priapulid transcriptomes/genomes need to be examined to ensure that the truncation is not an artifact caused during sequence assembly. The inability to detect more neuropeptides from these datasets could be attributed to query proteomes not enriched from neural tissue. Nonetheless, our analyses showcase the power of NP-HMMer by rapidly and reliably discovering neuropeptides in diverse protostomes. One limitation of the models generated here is that they are largely not suitable for discovering neuropeptides in non-bilaterians or deuterostomes. Therefore, we plan to expand NP-HMMer to include models for additional neuropeptides, including those that are predominantly found in non-protostomian invertebrates.

**Table 1:** Summary of neuropeptides identified in Priapulida and Rotifera.

|  | Priapulida | Rotifera |  |  |  |
| --- | --- | --- | --- | --- | --- |
| Precursors / Species | *Priapulus caudatus* | *Adineta ricciae* | *Brachionus plicatilis* | *Didymodactylos carnosus* | *Rotaria socialis* |
| ACP/APWGa |  |  | 1 |  |  |
| AKH |  |  | 1 |  |  |
| Agatoxin | 2 |  |  |  |  |
| Allatostatin-A |  | 1 | 1 | 2 | 2 |
| Allatostatin-B / Wamide | 1 | 1 |  | 1 |  |
| Allatostatin-C/CC/CCC | 2 | 1 |  |  |  |
| Bursicon alpha | 2 | 1 |  |  |  |
| Bursicon beta | 1 |  |  |  |  |
| Calcitonin | 1 |  |  |  |  |
| CAPA | 1 | 1 |  |  | 1 |
| CCAP | 1 |  |  | 2 | 1 |
| CCHa | 1 | 2 |  |  | 2 |
| CNMa | 1 |  |  |  |  |
| DH31 | 1 |  |  |  |  |
| DH44 | 1 |  |  |  |  |
| ETH | 1 |  |  |  |  |
| Eclosion Hormone | 3 |  |  |  |  |
| FMRFa |  | 1 | 1 | 2 | 2 |
| GPA2 | 2 |  |  |  | 2 |
| GPB5 | 1 |  |  |  |  |
| Insulin/IGF-like | 2 | 1 |  |  | 1 |
| ITP | 1 |  |  |  |  |
| Leucokinin |  |  |  | 2 | 1 |
| Neuropeptide-F | 1 | 5 | 3 | 1 | 3 |
| NUCB | 1 | 2 | 1 | 1 | 2 |
| Orcokinin-A/B | 1 | 2 |  | 1 | 2 |
| PDF |  | 1 |  |  |  |
| Phoenixin |  |  |  | 1 | 1 |
| Proctolin |  | 1 |  |  |  |
| PTTH | 1 |  |  |  |  |
| RYa/Luqin | 1 |  |  |  |  |
| Tachykinin | 1 |  |  |  |  |
| TRH | 1 |  |  |  |  |
| Vasopressin/Oxytocin |  | 2 | 1 |  | 2 |

**Figure 2:**
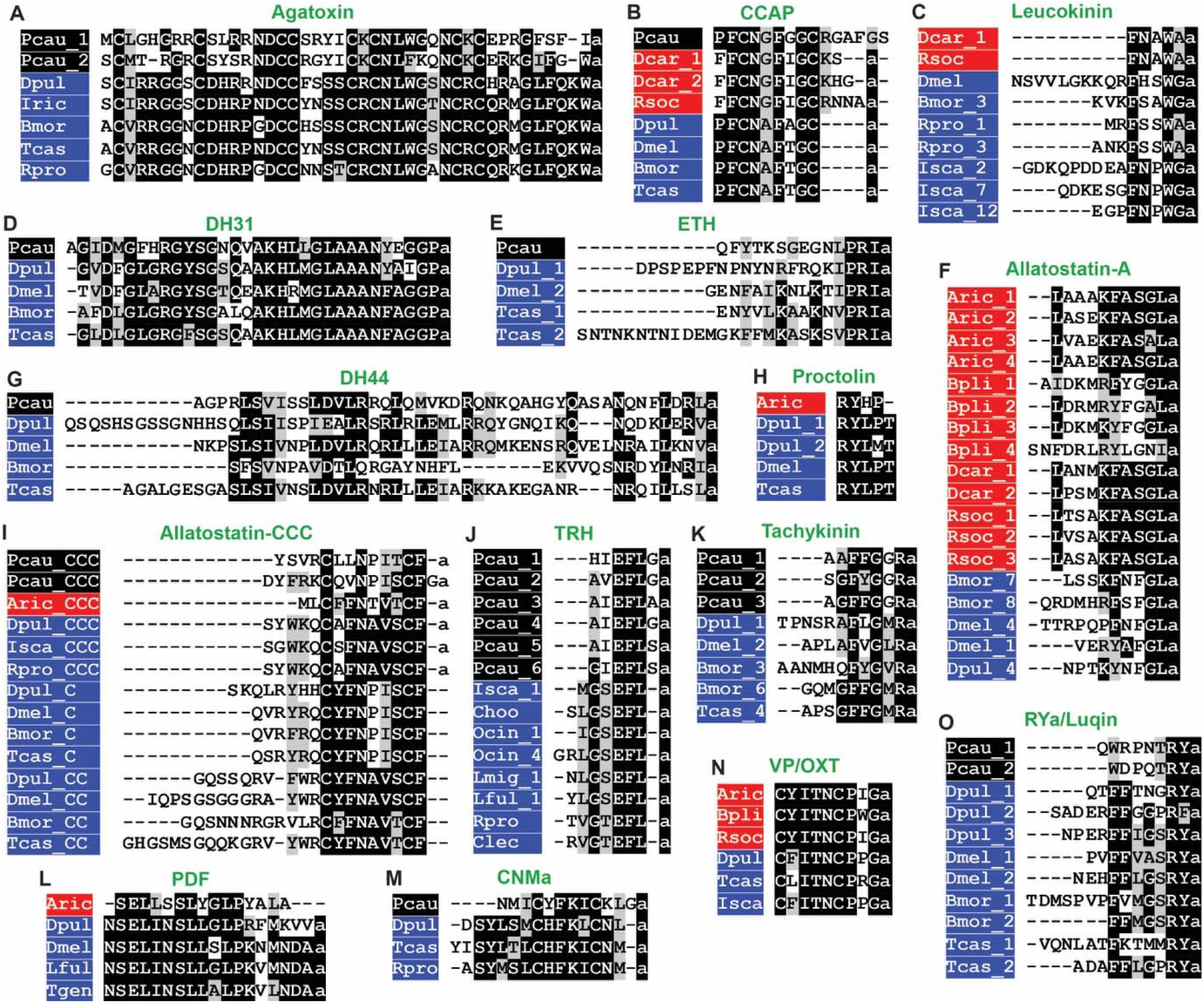
Multiple sequence alignments of neuropeptides discovered in Priapulida and Rotifera. Alignments of **(A)** agatoxin, **(B)** crustacean cardioactive peptide (CCAP), **(C)** leucokinin, **(D)** diuretic hormone 31 (DH31), **(E)** ecdysis-triggering hormone (ETH), **(F)** allatostatin-A, **(G)** diuretic hormone 44 (DH44), **(H)** proctolin, **(I)** allatostatin-CCC, **(J)** thyrotropin-releasing hormone (TRH), **(K)** tachykinin, **(L)** pigment-dispersing factor (PDF), **(M)** CNMamide (CNMa), **(N)** vasopressin/oxytocin (VP/OXT) and **(O)** RYamide (RYa)/Luqin mature peptides. Priapulid species are colored in black, rotifers in red, and arthropods in blue. Conserved residues are highlighted in black or gray. Species names: Pcau, *Priapulis caudatus*; Dcar, *Didymodactylos carnosus*; Rsoc, *Rotaria socialis*; Aric, *Adineta ricciae*; Bpli, *Brachionus plicatilis*; Dpul, *Daphnia pulex*; Iric, *Ixodes Ricinus*; Bmor, *Bombyx mori*; Tcas, *Tribolium castaneum*; Rpro, *Rhodnius prolixus*; Dmel, *Drosophila melanogaster*; Isca, *Ixodes scapularis*; Lful, *Ladona fulva*; Tgen, *Timema genevievae*; Choo, *Clitarchus hookeri*; Ocin, *Orchesella cincta*; Lmig, *Locusta migratoria*; Clec, *Cimex lectularius*.

**Figure 3:**
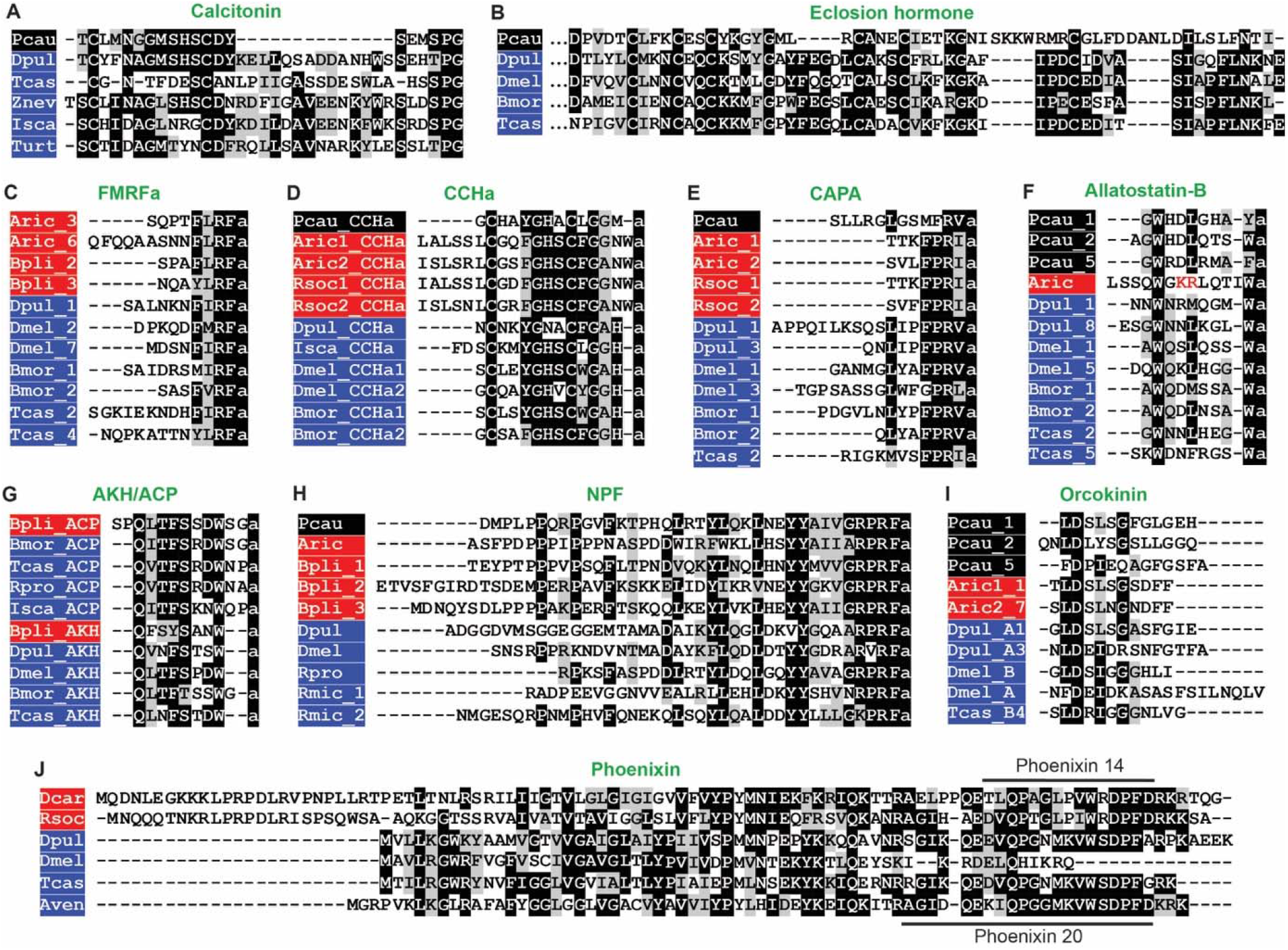
Multiple sequence alignments of neuropeptides discovered in Priapulida and Rotifera. Alignments of **(A)** calcitonin, **(B)** eclosion hormone, **(C)** FMRFamide (FMRFa), **(D)** CCHamide (CCHa), **(E)** CAPA, **(F)** allatostatin-B, **(G)** adipokinetic hormone (AKH) and AKH/corazonin-related peptide (ACP), **(H)** neuropeptide F (NPF), **(I)** orcokinin and **(J)** phoenixin neuropeptides. Only mature peptides are shown except for eclosion hormone (partial alignment) and phoenixin (complete neuropeptide precursor). A dibasic cleavage site (KR) in *Adineta ricciae* allatostatin-B is shown in red. These residues are likely cleaved to generate two peptides (LSSQWamide and LQTIWamide). Priapulid species are colored in black, rotifers in red, and arthropods in blue. Conserved residues are highlighted in black or gray. Species names: Pcau, *Priapulis caudatus*; Dcar, *Didymodactylos carnosus*; Rsoc, *Rotaria socialis*; Aric, *Adineta ricciae*; Bpli, *Brachionus plicatilis*; Dpul, *Daphnia pulex*; Bmor, *Bombyx mori*; Tcas, *Tribolium castaneum*; Rpro, *Rhodnius prolixus*; Dmel, *Drosophila melanogaster*; Isca, *Ixodes scapularis*; Rmic, *Rhipicephalus microplus*; Znev, *Zootermopsis nevadensis*; Turt, *Tetranychus urticae*; Aven, *Araneus_ventricosus*.

## Acknowledgments

We would like to thank Dr. Ismail Moghul and Dr. Maurice Elphick for the initial discussions and Dr. Theresa McKim for helpful feedback during the preparation of this manuscript. L.A.Y.G. was supported by a BBSRC fellowship (BB/W010305/1) and funding from the Royal Society (RG\R1\241397).

## Author contributions

Study conception: L.A.Y.G. and M.Z.; Computational analyses: L.A.Y.G., J.S. and W.Y.; Data analyses: M.B.A., L.A.Y.G. and M.Z.; Data visualization: all authors; Manuscript writing: M.Z.; Manuscript editing: all authors.

## Declaration of competing interest

We declare that we have no competing interests.

## Code availability

All the models, code, and files to automatically create the alignments are freely available on Github (https://github.com/Imnotabioinformatician/NP-HMMer)

---

**Table S1:** Summary of neuropeptides recovered using NP-HMMer in proteomes of model arthropods. Closely-related neuropeptides identified for a given model are included in brackets. Neuropeptides highlighted in yellow indicate sequences that were not part of the proteome used for analysis but were successfully recovered by NP-HMMer following their addition to the proteome. Phoenixin neuropeptide was identified for the first time in *Drosophila melanogaster*.

| Precursors / Species | *Drosophila melanogaster* | *Daphnia pulex* | *Tribolium castaneum* |
| --- | --- | --- | --- |
| ACP | 1 (1 AKH) | 1 (1 AKH) | 3 (2 AKH) |
| AKH | 1 | 1 | 3 (1 ACP) |
| Agatoxin | 0 | 1 | 1 |
| Allatostatin-A | 1 | 1 | 0 |
| Allatostatin-B | 1 | 1 | 2 (1 Tachykinin) |
| Allatostatin-C/CC/CCC | 2 (1 Allatostatin-CC) | 1 (1 Allatostatin-CCC) | 2 (1 Allatostatin-CC) |
| Allatotropin | 0 | 1 | 1 |
| Bursicon alpha | 1 | 2 (1 Bursicon beta) | 2 (1 Bursicon beta) |
| Bursicon beta | 1 | 2 (1 Bursicon alpha) | 2 (1 Bursicon alpha) |
| Calcitonin | 0 | 1 | 1 |
| CAPA | 1 | 1 | 2 (1 PBAN) |
| CCAP | 1 | 1 | 1 |
| CCHa-1 | 2 (1 CCHa-2) | 1 (1 CCHa) | 2 (1 CCHa-2) |
| CCHa-2 | 2 (1 CCHa-1) | 1 (1 CCHa) | 2 (1 CCHa-1) |
| CNMa | 1 | 1 | 1 |
| Corazonin | 1 | 1 | 1 (1 ACP) |
| DH31 | 1 | 1 | 1 |
| DH44 | 1 | 1 | 2 |
| ETH | 1 | 1 | 1 |
| Eclosion Hormone | 1 | 2 | 2 |
| Elevenin | 0 | 1 | 1 |
| FMRFa | 1 | 1 | 1 |
| GPA2 | 1 | 1 | 1 |
| GPB5 | 1 | 1 | 1 |
| Insulin/IGF-like | 6 | 3 | 4 |
| ITP | 1 | 3 | 1 |
| Leucokinin | 1 | 0 | 0 |
| Myosuppressin | 1 | 1 | 1 |
| Natalisin | 1 | 1 | 2 (1 Tachykinin) |
| Neuroparsin | 0 | 1 | 1 |
| Neuropeptide-F | 1 | 1 | 0 |
| NUCB | 1 | 1 | 1 |
| Orcokinin-A/B | 1 (1 Orcokinin-A) | 1 (1 Orcokinin-A) | 1 (1 Orcokinin-B) |
| PDF | 1 | 1 | 1 |
| Phoenixin | 1 | 1 | 1 |
| Proctolin | 1 | 1 | 1 |
| PTTH | 1 | 0 | 1 |
| RYa/Luqin | 1 | 1 | 1 |
| SIFa | 1 | 1 | 1 |
| sNPF | 1 | 1 | 1 |
| Sulfakinin | 1 | 1 | 1 |
| Tachykinin | 1 | 2 (1 Natalisin) | 2 (1 Allatostatin-B) |
| TRH | 0 | 1 | 0 |
| Trissin | 1 | 0 | 1 |
| Vasopressin/Oxytocin | 0 | 1 | 1 |

## Notes

https://github.com/Imnotabioinformatician/NP-HMMer

### Competing Interest Statement

The authors have declared no competing interest.

