## Supplementary material for "Proteome-wide neuropeptide identification using NeuroPeptide-HMMer (NP-HMMer)": Figure S1_Drosophila_neuropeptidome.docx

**Figure S1:** *Drosophila melanogaster* neuropeptides recovered using NP-HMMer.

>AKH | Dmel3465

MNPKSEVLIAAVLFMLLACVQCQLTFSPDWGKRSVGGAGPGTFFETQQGNCKTSNEMLLEIFRFVQSQAQLFLDCKHRE

>Allatostatin-A | Dmel7638

MNSLHAHLLLLAVCCVGYIASSPVIGQDQRSGDSDADVLLAADEMADNGGDNIDKRVERYAFGLGRRAYMYTNGGPGMKRLPVYNFGLGKRSRPYSFGLGKRSDYDYDQDNEIDYRVPPANYLAAERAVRPGRQNKRTTRPQPFNFGLGRR

>Allatostatin-B | Dmel11489

MAHTKTRRTYGFLMVLLILGSACGNLVASGSAGSPPSNEPGGGGLSEQVVLDQLSESDLYGNNKRAWQSLQSSWGKRSSSGDVSDPDIYMTGHFVPLVITDGTNTIDWDTFERLASGQSAQQQQQQPLQQQSQSGEDFDDLAGEPDVEKRAWKSMNVAWGKRRQAQGWNKFRGAWGKREPTWNNLKGMWGKRDQWQKLHGGWGKRSQLPSN

>Allatostatin-C | Dmel9436

MMKFVQILLCYGLLLTLFFALSEARPSGAETGPDSDGLDGQDAEDVRGAYGGGYDMPAQAIYPNIPMDRLQMLFAQYRPTSYSAYLRSPTYGNVNELYRLPESKRQVRYRQCYFNPISCFRK

>Allatostatin-CC | Dmel13714

MHQPPGRQTARRRRSCTSLAGKEGTPLCRTYHLPAMLIILLVLIQNFELHMCRQLMVYPGADKRSPDKLLTIGGSAAGEVTLPEANTPADDKRAGGSRSAPSQPEEIFSAPADEGYDEYPMVVPKRAALLLDRLMVALHHALEQERSEQRIGEFFGDRNILSGKFGDSHNGMEHHQAREDGMYSDDDAGTLLDYDFKDLNQINRATGETLMPKSFQRRAGADRSGTSTHSGSPAGSRRIQPSGSGGGRAYWRCYFNAVSCF

>Bursicon alpha | Dmel7883

MLRHLLRHENNKVFVLILLYCVLVSILKLCTAQPDSSVAATDNDITHLGDDCQVTPVIHVLQYPGCVPKPIPSFACVGRCASYIQVSGSKIWQMERSCMCCQESGEREAAVSLFCPKVKPGERKFKKVLTKAPLECMCRPCTSIEESGIIPQEIAGYSDEGPLNNHFRRIALQ

>Bursicon beta | Dmel9254

MHVQELLFVAAILVPQCLRALRYSQGTGDENCETLKSEIHLIKEEFDELGRMQRTCNADVIVNKCEGLCNSQVQPSVITPTGFLKECYCCRESFLKEKVITLTHCYDPDGTRLTSPEMGSMDIRLREPTECKCFKCGDFTR

>CAPA | Dmel6699

MKSMLVHIVLVIFIIAEFSTAETDHDKNRRGANMGLYAFPRVGRSDPSLANSLRDGLEAGVLDGIYGDASQEDYNEADFQKKASGLVAFPRVGRGDAELRKWAHLLALQQVLDKRTGPSASSGLWFGPRLGKRSVDAKSFADISKGQKELN

>CCAP | Dmel7809

MRTSMRISLRLLALLACAICSQASLERENNEGTNMANHKLSGVIQWKYEKRPFCNAFTGCGRKRTYPSYPPFSLFKRNEVEEKPYNNEYLSEGLSDLIDINAEPAVENVQKQIMSQAKIFEAIKEASKEIFRQKNKQKMLQNEKEMQQLEERESK

>CCHa-1 | Dmel4197

MWYSKCSWTLVVLVALFALVTGSCLEYGHSCWGAHGKRSGGKAVIDAKQHPLPNSYGLDSVVEQLYNNNNNNQNNQDDDNNDDDSNRNTNANSANNIPLAAPAIISRRESEDRRIGGLKWAQLMRQHRYQLRQLQDQQQQGRGRGGQGQYDAAAESWRKLQQALQAQIDADNENYSGYELTK

>CCHa-2 | Dmel6139

MKSTISLLLVVICTVVLAAQQSQAKKGCQAYGHVCYGGHGKRSLSPGSGSGTGVGGGMGEAASGGQEPDYVRPNGLLPMMAPNEQVPLEGDFNDYPARQVLYKIMKSWFNRPRRPASRLGELDYPLANSAELNGVN

>CNMa | Dmel12508

MSALSAPTTCGCSPVHWAIVIVLLSVAIGPGDAMARPARNTQLLFSELLGGGNDDNNYYGDQLKYQQQQQQQQEQKQQRVPAFARKWPSLRDLLLTVDYDDFGVTQESEEQVAPSSRLLARLHRLGDNGGGEELRYNVVNELTNMPSKKVMPGHPLKDHNTKKNVQFRKQYMSPCHFKICNMGRKRNAGFNSY

>Corazonin | Dmel3981

MLRLLLLPLFLFTLSMCMGQTFQYSRGWTNGKRSFNAASPLLANGHLHRASELGLTDLYDLQDWSSDRRLERCLSQLQRSLIARNCVPGSDFNANRVDPDPENSAHPRLSNSNGENVLYSSANIPNRHRQSNELLEELSAAGGASAEPNVFGKH

>DH31 | Dmel9661

MTNRCACFALAFLLFCLLAISSIEAAPMPSQSNGGYGGAGYNELEEVPDDLLMELMTRFGRTIIRARNDLENSKRTVDFGLARGYSGTQEAKHRMGLAAANFAGGPGRRRRSETDV

>DH44 | Dmel8725

MMKATAWFCPVLLTLLCATRLVCTAQRGAVGAGGAAGGSGAAAGGAEVGGSGRTNGYPLDYPDGTRNSQDDFLLAKRNKPSLSIVNPLDVLRQRLLLEIARRQMKENSRQVELNRAILKNVGKRVVLRGGGGGGGSGAGGLAPKVSRRYRQQWPVERELERERQRERERERDAVREEQLDRQQLLPWKHFPSQLWSYGWALSPYKESSQLQFADSQQSASTGPQSQALPKQLQLLSYAKKPLDVAGMSLARHRVSGNEANETNHENDDGNGASKNPARYVDDGDNEGEDSYNDVGTEGVGLGLGMGVGLGLERFEVLEDKPNWANEEPNELVVVNANDRVPWSFPYRFHKSQHNVN

>ETH | Dmel12663

MRIITVLSVSLLVGLVAISQADDSSPGFFLKITKNVPRLGKRGENFAIKNLKTIPRIGRSEHSSVTPLLAWLWDLETSPSKRRLPAGESPAKEQELNVVQPVNSNTLLELLDNNAIPSEQVKFVHWKDFDRALQADADLYSKVIQLGRRPDQHLKQTLSFGSFVPIFGDEQNPDFMMYKNNEDQELYGGGNRYDRQFLKYNIL

>Eclosion Hormone | Dmel3664

MNCKPLILCTFVAVAMCLVHFGNALPAISHYTHKRFDSMGGIDFVQVCLNNCVQCKTMLGDYFQGQTCALSCLKFKGKAIPDCEDIASIAPFLNALE

>FMRFa | Dmel2853

MGIALMFLLALYQMQSAIHSEIIDTPNYAGNSLQDADSEVSPSQDNDLVDALLGNDQTERAELEFRHPISVIGIDYSKNAVVLHFQKHGRKPRYKYDPELEAKRRSVQDNFMHFGKRQAEQLPPEGSYAGSDELEGMAKRAAMDRYGRDPKQDFMRFGRDPKQDFMRFGRDPKQDFMRFGRDPKQDFMRFGRDPKQDFMRFGRTPAEDFMRFGRTPAEDFMRFGRSDNFMRFGRSPHEELRSPKQDFMRFGRPDNFMRFGRSAPQDFVRSGKMDSNFIRFGKSLKPAAPESKPVKSNQGNPGERSPVDKAMTELFKKQELQDQQVKNGAQATTTQDGSVEQDQFFGQ

>GPA2 | Dmel651

MGSSQLLVLICCIPWLCDSNSMGKDAWLRPGCHKVGNTRKITIPDCVEFTITTNACRGFCESFSVPSIPMMGSSLSVLFKPPKPVVSVGQCCNMMKSEEIQRRVLCIEGIRNVTFNSALSCSCYHCKKD

>GPB5 | Dmel4193

MLADSATLTLAIFVGTSVVLVSVSSSSLSEIKPMNNGHIVTPLGCHRRVYTYKVTQSDLQGHECWDYVSVWSCWGRCDSSEISDWKFPYKRSFHPVCVHAQRQLVVAILKNCHPKAEDSVSKYQYMEAVNCHCQTCSTQDTSCEAPANNEMAGGSRAIMVGADTKNLDY

>Insulin-like_Ilp1 | Dmel11015

MFSQHNGAAVHGLRLQSLLIAAMLTAAMAMVTPTGSGHQLLPPGNHKLCGPALSDAMDVVCPHGFNTLPRKRESLLGNSDDDEDTEQEVQDDSSMWQTLDGAGYSFSPLLTNLYGSEVLIKMRRHRRHLTGGVYDECCVKTCSYLELAIYCLPK

>Insulin-like_Ilp2 | Dmel11016

MSKPLSFISMVAVILLASSTVKLAQGTLCSEKLNEVLSMVCEEYNPVIPHKRAMPGADSDLDALNPLQFVQEFEEEDNSISEPLRSALFPGSYLGGVLNSLAEVRRRTRQRQGIVERCCKKSCDMKALREYCSVVRN

>Insulin-like_Ilp3 | Dmel11017

MGIEMRCQDRRILLPSLLLLILMIGGVQATMKLCGRKLPETLSKLCVYGFNAMTKRTLDPVNFNQIDGFEDRSLLERLLSDSSVQMLKTRRLRDGVFDECCLKSCTMDEVLRYCAAKPRT

>Insulin-like_Ilp4 | Dmel11018

MSLIRLGLALLLLLATVSQLLQPVQGRRKMCGEALIQALDVICVNGFTRRVRRSSASKDARVRDLIRKLQQPDEDIEQETETGRLKQKHTDADTEKGVPPAVGSGRKLRRHRRRIAHECCKEGCTYDDILDYCA

>Insulin-like_Ilp5 | Dmel5116

MMFRSVIPVLLFLIPLLLSAQAANSLRACGPALMDMLRVACPNGFNSMFAKRGTLGLFDYEDHLADLDSSESHHMNSLSSIRRDFRGVVDSCCRKSCSFSTLRAYCDS

>Insulin-like_Ilp7 | Dmel13425

MTRMIIQNSGSWTLCGAVLLFVLPLIPTPEALQHTEEGLEMLFRERSQSDWENVWHQETHSRCRDKLVRQLYWACEKDIYRLTRRNKKRTGNDEAWIKKTTTEPDGSTWLHVNYANMFLRSRRSDGNTPSISNECCTKAGCTWEEYAEYCPSNKRRNHY

>ITP | Dmel3715

MCSRNIKISVVLFLVLIPIFAALPHNHNLSKRSNFFDLECKGIFNKTMFFRLDRICEDCYQLFRETSIHRLCKANCFVHETFGDCLKVLLIDDEEISQLQHYLKVINGSPYPFHKPIYH

>Leucokinin | Dmel3471

MAKIVLCMVLLAFGRQVYGASLVPAPISEQDPELATCELQLSKYRRFILQAILSFEDVCDAYSSRPGGQDSDSEGWPFRHYAPPPTSQRGEIWAFFRLLMAQFGDKEFSPIIRDAVIERCRIKSQLQRDEKRNSVVLGKKQRFHSWGGKRSPEPPILPDY

>Myosupressin | Dmel3463

MSFAQFFVACCLAIVLLAVSNTRAAVQGPPLCQSGIVEEMPPHIRKVCQALENSDQLTSALKSYINNEASALVANSDDLLKNYNKRTDVDHVFLRFGKRR

>Natalisin | Dmel1970

MRLTLAWLSLCLAIYCGGGHGHGNVVLSLPPSLIATATKAALSHQRQQKQQQQHQKKDARVLFDSPADALRDMMHNGNGNGNGPMDSGKFSLSDVEQPAAQRSEDFNRNAYDLGARQSAPQEIAMGMELGMGLGLGPNNYRTTPPHRYWGQRCQGRSGGSGTSKCPQEYYRTMLAARNKEALSRLHMQLSSMQDSDSGASSSSDSEEEHVDDEEQSNNEVFMLLTGEQDLMKFLHWAMQVLYPIERPLGNLSDGAAENYYPGMFLWKKLNLSGHLEPPLIVDEPQYVLVRREKLFDGYQFGEDMSKENDPFIPPRGRKHSGSLDLDALMNRYEPFVPNRGKRDKVKDLFKYDDLFYPHRGKKHRNLFQVDDPFFATRGKKLQLRDLYNADDPFVPNRGKRHLTASAGKLGETMAGGGKWPDDSNNYWPLRMSTHKINGYDQSVRPSLSVEDAAASLASWRLPANRLHSTRSMSADLRQQLLLPHVRFIGNPNMRQQQQQQQQHQVKTSSWQAEERLRRSILAPGESNDAHETQLTLSHPANPHLVTDTDNLNI

>Neuropeptide-F | Dmel8198

MCQTMRCILVACVALALLAAGCRVEASNSRPPRKNDVNTMADAYKFLQDLDTYYGDRARVRFGKRGSLMDILRNHEMDNINLGKNANNGGEFARGFNEEEIF

>NUCB | Dmel11523

MVQNVALLGLALIAISASIVALPVTQNKKDHKEAAESSTPATADVETALEYERYLREVVEALEADPEFRKKLDKAPEADIRSGKIAQELDYVNHHVRTKLDEIKRREVERLRELANQAYELSNDIDRKHLKVSQHLDHDNEHTFEIEDLRKLIQKTSDDLAEADRKRRGEFKEYEMQKEFEREAQKKEMDEESRKKFETELKEKEEKHKDHEKLHHPGNKAQLEDVWEKQDHMDKNDFDPKTFFSIHDVDSNGYWDEAEVKALFVKELDKVYQSDLPEDDMRERAEEMERMREHYFQETDMNHDGLISIDEFMVQTNKEEFQKDPEWETIDRQQQYTHEEYLEYERRRQEEVQRLIAQGQLPPHPNMPQGYYAAPPPGGVAYQQAPPGAQLHYQHPDQVHAQQQQQYAQQQQQYAQQYQQQQYGNGQQPVQLQPNQVYQHAGQIPQQQQPVYQNQPVYQQQQPVYQQQQPVQQQQKPVQQPVQQQQQPVQQQQQPVQQQQQTVQQQQPVQQQQQTVQQQQPVQQQQQTAQQQPVAQQQIHNQSPPPVLNQQVPVQQQQKQHQESLNQQH

>Orcokinin-A | Dmel12751

MNLYVLLAVVSVFLNFIHAAPGVDISNDELLDGKYLCEAGSKKYDGPFIVRLISAANGQTVVCYECSQSEFKTKYSVKQCAAGKIGSGHHRDLVPYLVRMDPLYKDTWSSKLKRNFDEIDKASASFSILNQLV

>PDF | Dmel2661

MARYTYLVALVLLAICCQWGYCGAMAMPDEERYVRKEYNRDLLDWFNNVGVGQFSPGQVATLCRYPLILENSLGPSVPIRKRNSELINSLLSLPKNMNDAGK

>Phoenixin | Dmel660

MAVLRGWRFVGFVSCIVGAVGLTLYPVIVDPMVNTEKYKTLQEYSKIKRDELQHIKRQ

>Proctolin | Dmel9730

MGVPRSHGTGIGCGSGHRWLLVWMTVLLLVVPPHLVDGRYLPTRSHGDDLDKLRELMLQILELSNEDPQQQQQQQQQQQHPQLRLHNEATGGSSSSSNINNPRVSNGNSNAAWLQKLSAMGALDELGGDGARFGPNYGRY

>PTTH | Dmel13757

MDIKVWRLLGQSCRRSTALVRHRSSMMPWLPISIMAMALLSLTNKGIASQYASDEGLDEMVGLRSLEHRAEEQQPDRSTSKMLSALFGFSPSTPHPTEMAMVMPHQLPPMYYNDFYEDLVTTKRNDVHSAGCDCKVTNELVDLGGLHFPRFLMNAVCESGAGRDLAKCSHGSNCRPLEYKVKVLAQTSQSDHPYSWMNKDQPWQFKTVTVTAGCFCTK

>RYamide/Luqin | Dmel2009

MNECVNKLLHLKFLFYFILGIQKRPVFFVASRYGRSTTYDESLKSRRIFIVPRNEHFFLGSRYGKRSGKYLCLSREINKLIVRKRLRNNDKERTPTLSFITKHFLMRNT

>SIFa | Dmel4360

MALRFTLTLLLVTILVAAILLGSSEAAYRKPPFNGSIFGKRNSLDYDSAKMSAVCEVAMEACPMWFPQNDSK

>sNPF | Dmel9022

MFHLKRELSQGCALALICLVSLQMQQPAQAEVSSAQGTPLSNLYDNLLQREYAGPVVFPNHQVERKAQRSPSLRLRFGRSDPDMLNSIVEKRWFGDVNQKPIRSPSLRLRFGRRDPSLPQMRRTAYDDLLERELTLNSQQQQQQLGTEPDSDLGADYDGLYERVVRKPQRLRWGRSVPQFEANNADNEQIERSQWYNSLLNSDKMRRMLVALQQQYEIPENVASYANDEDTDTDLNNDTSEFQREVRKPMRLRWGRSTGKAPSEQKHTPEETSSIPPKTQN

>Sulfakinin | Dmel2817

MGPRSCTHFATLFMPLWALAFCFLVVLPIPAQTTSLQNAKDDRRLQELESKIGGEIDQPIANLVGPSFSLFGDRRNQKTMSFGRRVPLISRPIIPIELDLLMDNDDERTKAKRFDDYGHMRFGKRGGDDQFDDYGHMRFGR

>Tachykinin | Dmel8553

MRPLSGLIALALLLLLLLTAPSSAADTETESSGSPLTPGAEEPRRVVKRAPTSSFIGMRGKKDEEHDTSEGNWLGSGPDPLDYADEEADSSYAENGRRLKKAPLAFVGLRGKKFIPINNRLSDVLQSLEEERLRDSLLQDFFDRVAGRDGSAVGKRAPTGFTGMRGKRPALLAGDDDAEADEATELQQKRAPVNSFVGMRGKKDVSHQHYKRAALSDSYDLRGKQQRFADFNSKFVAVRGKKSDLEGNGVGIGDDHEQALVHPWLYLWGEKRAPNGFLGMRGKRPALFE

>Trissin | Dmel4230

MTKTTMHWLAHFQIILLCIWLMCPPSSQAIKCDTCGKECASACGTKHFRTCCFNYLRKRSDPDALRQSSNRRLIDFILLQGRALFTQELRERRHNGTLMDLGLNTYYP
