## Supplementary material for "Proteome-wide neuropeptide identification using NeuroPeptide-HMMer (NP-HMMer)": Figure S2_Daphnia_neuropeptidome.docx

**Figure S2**: *Daphnia pulex* neuropeptides recovered using NP-HMMer.

>AKH | Dpul28768

MANHRILILTLLMIGLASAQVNFSTSWGKRSPSTSTKAAEPPSAPSYRQNFHSKKVEPGTLETLPNNQHLPESFDTVSSTIYDDAEEQRISISLPSPCLSILKSLLLVNQIVEFKNSPLDGRMHRFKIENLFPLPNRTCRLYIRR

>Agatoxin | Dpul12004

MTNEQQSAGLNYFTDNAEPLMENAQKRSCIRRGGSCDHRRNDCCFSSSCRCNLWGSNCRCHRAGLFQKWG

>Allatostatin-A | Dpul29718

MLFLSTALFLLVALLQSSQCTDVNNDSSVIEAVGAGGGDSQAKAESLASAEDKLSALTADGMATRARYFKSFGGNPTGDPNLNIYSFGLGKRTSRSYSINPYSFGLGKRGGNAKSYPQQIPYSFGLGKRNPTKYNFGLGKRPDRFGFGLGKRNLKDDDLDQWLNDEEYSQFDDTEEEIEGREDDSQEIMDVNKRSQMAAGQQQQQQQQAFFPSHLQSAFYGGPFMSNLARANTHALGKSRTFNQQAATHDLPNLGKRLPVYNFGLGK

>Allatostatin-B | Dpul745

MQFWQCPLLLMVSLIAAINTQQTPSQRLEPNQVAGLVELHQQLEQPREHQQQQQHHQQQPEQPAQADNKDYAPSPAVLLQLTPSDWKNTQDKRNNWNRMQGMWGKRSQQQSDESALSEMTPPRMVKRAWSDLSQQGWGKRSWTQLHGVWGKRRWDQLHGAWGKRTPDQLEDDSKAEQPENSQEDEDQQSSSEVEREEANDSDDTVENSKRSGWNKMQGVWGKRSSSSSKTNSGPAIEGMGNNDLLLLISGTGDQLYQQREDDQAKANVEDAGNEESVDKRGWNQLQGVWGKRALSAMAAGYKRNWNNLRGAWGKREIPAAIAKGMEWSRKRESGWNNLKGLWG

>Allatostatin-CCC | Dpul8516

MMAKISAVVPVAILLYLAASGAAKSTDREETESTDFGQDIEVLGAVPDDGSVETALLNYLFAKQIVARLRTNANPQDLMRKRSYWKQCAFNAVSCFGK

>Allatotropin | Dpul10065

MKGKGAFLMVLAGWGLIGLMILTTAVEAAPHPADYTSSSVNNQRDFRSRRGFKTVGLATARGFGKRAPSLSNFNSFQDAAEQMMQQQEENPNSDPDVFPVDWLVNYLQNKPDVIRYMVEHLLDHNGDGQVTSQEMMTSLQQQRED

>Bursicon alpha | Dpul22408

MFLFVGLVSVIRADECQLTPVIHVLQYPGCIPKPIPSFACTGKCTSYVQVSGSKLWQTERSCMCCQESGEREATVSLLCPKAAPGEPKLRRVVTRAPVDCMCRPCTALEESAVMPQEIARFLDDGSFPFKL

>Bursicon beta | Dpul29409

MFWYVLIILLGAGSSRETLASKTNLMSGTCETLPSTIHITKEEYTDGGILSRTCEGDIGVAKCEGSCSSQVQPSVVHPSGFLKECMCCRESFLRERVVTLTHCYDANGNRLTGKSSSLDVKMREPADCKCFRCGDSAE

>Calcitonin | Dpul1069

MNDNMYADIELIEDYPMKPVTMFDILKRVLAQKPAPPEPTELIKTPLIRKRTCYFNAGMSHSCDYKELLQSADDANHWSSEHTPGKRRKRSSRGRAPAAASTSVPEKAAAGVVREIKKEEKELDITVDAMRVVGTSHTSRILCFSFVEKGAAICAHTQLEFSIAARELRYHIWCRSQSVTMLEALTRQSTALALYIEPCC

>CAPA | Dpul16615

MRIAIIHSLVLVVIYLAYSDAAPPQILKSQSLIPFPRVGRSRSSFIANAVGSARSGGAGNPMMMGSGGNANGKSPNNWMMNNADIKRHLIPFPRVGKRQNLIPFPRVGRAGYYQPGFFPSTDDEEGQASIQQQFALSSEESQASAPSSFLMSGSDILGALNNGRSNSDERTAVFIPRRWMTNSQESEE

>CCAP | Dpul16169

MTRPLFYSLLMLAWMIISLYISASSSQPLKNNQNDSDSAEEIEQWSFKEKRPFCNAFAGCGRKRSMIKDTKHPPYEKAASNHPRLPNADTKLLDKLFAKIQHQRANFVQLDDPEYY

>CCHa | Dpul17626

MHIFFYVIHVTAMLAIVSGNCNKYGNACFGAHGKRSDFKRTSAVDLSDQIWPVAANWNPTRPDEPIQERRQMKPLPALQLESVLVYNDIPRSAEHSRYLNQEDYNN

>CNMa | Dpul20247

MLPFERLNPRFWLGPEYRQALRQIAQMREEDPQIFNGLNVYRAADGFIAQGSNAMDRTDIGVRLSPARERLQVANPDAQSNMGKRDSYLSMCHFKLCNLGRKRRISQGGHTDLVTSDLNAEK

>Corazonin | Dpul23911

MSRTRVRLNLCHPKLFESRSPTEYQSKTAQSQRYSRGWTNGRKRSDPSFVQQQQWIQRNGHPMVIPAEFRSNSFEDWSRYRINGEKVNEDGDSWLVHVSHCAKLATSLGSVLKNKDAKSDDNPLIDVIH

>DH31 | Dpul28001

MSRFVMTIFFLLVACLALIVPGSAAPPRRPMLVDLDDPDSVMEVITRLERSLLRNSDYEHQKRGVDFGLGRGYSGSQAAKHLMGLAAANYAIGPGRKRRDTTESTPEDVKTGAIN

>ETH | Dpul13157

MYRELIMIKGLFLTWLLLASALSDPSPEPFNPNYNRFRQKIPRIGRRGEGIIAEYMNSESFPHEGSLSNFFLKASKAVPRLGRRKDISTESGRAAMVGEEPFGRISNEIPIMNQKQDLWPNMNINELTGALNKELNYPGPRIPKDLQDNYIQDLIHSWINQYENLNEN

>Eclosion Hormone | Dpul8527

MYGEYFEGQRCAEFCLASYKPSQAGSVSGGGGGSGWAPMPDCNEPETVDQFLKLSLMPQSDGPSVQQLIDDSDSDGGGGYMSPISQALMGSAYSGRYAGRKPSASAQQNTLYSSRTGAEFKGHNQLLKNRKWKKRINGSSKLLGGSPSMPHLVGY

>Eclosion Hormone | Dpul8721 (partial)

MTVDVAQPADTLYLCMKNCEQCKSMYGAYFEGDLCAKSCFRLKGAFIPDCIDVASIGQFLNKNE

>Elevenin | Dpul15101

MMRFNSMSSSSTILLLFAGIVIFAAVHVHSRDLDCRRFVFAPMCRGATIKRSFIPNIGVNQDDVHELLLDYPKQQVMDAETQPLLIPILIDRRIFEARSAANRHQPQLDAED

>FMRFa | Dpul27178

MNGLRFLMLILGLMMIVGQVRPDDETSEEDVDDENFLSGSNAAAAAGSSDSAEEDDDNNNNKGGIELYLSAMKDLYRQSKRHEAEPLVSSRSGASAPIAALYRRSALNKNFIRFGRSGGGVMKTDVSRQMQPIRLMGLNESDSGKDEERSFHPARPSRSLRSNFIRFGRSFFPTSNRWGETVRRSARRTPSAGRQMMSLVDLPASSSSSSRR

>GPA2 | Dpul28427

MVSTLYFWLVLAVLLTASNGEPRNQPKGSISVSTRSSTSRGELSGCHQVGHTRRVTIPDCVSFMITTNACRGFCESWSVPSSWEALLKNPEKVITSVGQCCNIMASEDVTVRVMCLGGPRDFTFKSAKTCSCFTCKKD

>GPB5 | Dpul929

MDATPTCFRRPYTFKVYQEDSEGRSCWDVVTVTSCWGRCSSNEIADWRFPFKRSQHPVCQHDSILPRAIILRNCDPEVNPGTELYMALDAVTCRCELCHTDTTNCEGPHYDRRITTHRLN

>Insulin-like | Dpul1320

MAHLTVGRCWMACIILLALATFTLARPPQENQPMTIRFCGRDLIRAIDEVCVAIKSPAAFIDPVALNQSSSLADSESADTQLRKWTKLEDEKSDSLLRQCCVIGCTDNDLTTFCQIDRNQIMGAAMHLRSQEWDMDWLYEFLPRYGDVPTNA

>Insulin-like | Dpul8939

MIIPSTVGRCWMALLLVLATLAFLTQSSPIHKRNVSGKNKWCGQEFSNALRAVCAHYKSYWPVTPAEPTEVGNDLKPQKELENRRSPTTNECCLVGCTQKDLESFCETVRNQTTVELPKSSDLSAANVMEWNSQPSEPLSTTETPSDSTPEIFYNESFEQDALTILPTYRTPQYSNLQFL

>Insulin-like | Dpul18367

MAYSTSFGSLVITICLLYSLTQLSAHTLQNPVNLGLLKDEETQARNYFADFLRVLYAPQAPKDDVSLSAGSSNDMPNEIFPMISSDEDDHEEENYDKIRRKEFPVDGVLLPILQKRNARYCGSYLADALRMACSKSSYLPLFGKRSTLPGKSLGLTTTAATPNAELGSWPFINDDKAHSILSNHHLFHRYTRGVHDECCVKGCTFKELTSYCTRPN

>ITP | Dpul14793

MDSGRQSHSSCSIRTLLGLTTLLVVLLAILPYYPTSAMSALSSDHHSSCKGLYDKSIFYRLNRICHDCFSLYRSPELHTLCRSECFTTPFFKACLKVLLMGDQDPDSSEMIDKIGR

>ITP | Dpul23863

MSALSSGHHSLSKRSFFDINCKGLYDKSIFARLDRICQDCYSLYREPELHTLCRKNCFTTNYFKGCLDALLINDEKDIQRVMKDISIIHQIPI

>ITP | Dpul26149

MASLILAEQPPGISIFPHPLSKHSFTEISKCDGVYDMNIYAQFNQICLNCYNLYRQPEIYRGCRKECFTSEYFGGCLAVLQITDKKDKILEDLSIIHRQMESPKEN

>Natalisin | Dpul6511

MELIKIIFVLASGWATLAIAGNTDQDMFWAARGKKASIEGNWPDSVVMEPFVAAVYDKRDGTFWAARGKKYAADGGDGVPFWATRGKKGDLEIPFWAARGKRIPQSEMNETEEEGNRREKRSAGRDTSIIHSGQRFRNRPARPASQAAEPFWAARGKKNSNVSKFPLL

>Neuroparsin | Dpul5478 (partial)

CVYGIVKDYCGRDICAKGPGHRCGGKWNSLGICGEGLFCSCNRCGGCSLNTIECFNLTCI

>Neuropeptide-F | Dpul27734 (partial)

MSSSNNSIQQFLPRSCSLAALVFLVVMAVLAVCVTTTKADGGDVMSGGEGGEMTAMADAIKYLQGLDKVYGQAARPR

>NUCB | Dpul13550

MELEMQQETGLEYNRYLQEVVQLLESDPDFRQKLEKSDPEDIRTGKVAKELEYVNHHVRNKLDELKRQEMERLRHLAMEEYERARGLGIPHDGRLKIPGHLDHKSPSFESEDLRKLIVQTSKDLEEADKQRKEEFKEYEMQKEFEHQNKLKGLDEEKRKTEEKEWEEAQAKHKQHPKMHHPGSKQQLEEVWEEQDHLSPDSFNPKTFFALHDLDGNGYWDPDEVKALFSKELDKAYDPNAPEDDMAERYEEMERMREHVFNETDTNRDYLISFPEFLEQTRKQEFERDPGWNTIDEQPVYSQQEYMEFEHQRQMEIQRLIDQGMLPPHPGMMPNHPRMPNVPYGVAPGQYAGQQPYYPSPVAQGKPITPEEAIRMQQQQYAQQPQFAPQQQYYGQPQFAQQQQYAGHPQQQFAANPQYAAQPQQPQFAQPEQQNFQQAAQQQQQNFQQASQQQQPIQQVPQQQPIAQQPVQQQQIPAQQQPLVQPQVPVQQQAPIQSQPVALPPAPQQIPAAKTV

>Orcokinin-A | Dpul29071

MNCLKFRLVAVAILIFNVVTALNYQSEEAVGVEHERDRDSLGGGHILRGLDSIGESNLLRAIYREKPRDFLRINRGLDSLSGASFGIEKRLDSLTGLGFGSQKRNLDEIDRSNFGTFAKRNLDEIDRSDFGRFVKKRETMEAESSQQQH

>PDF | Dpul13704

MHQLSAKLSHLSIALFVLLVSFATDAQSAPPSISSNNRPEAQMSIQEMEKFLEGLTRYLHRQHLDLPKVHQQSQEEQPGSYEADAIDRSGDMSAPTETERSSSSSSSELANHSLLSHPRPPMANKWPWSLSHLERIEDDPDFKERQQPYAKRNSELINSLLGLPRFMKVVG

>Phoenixin | Dpul22605

MVLLKGWKYAAMVGTVVGAIGLAIYPIIVSPMMNPEPYKKQQAVNRSGIKQEEVQPGNMKVWSDPFARPKAEEK

>Proctolin | Dpul22405

MLKSTSLKALVTLLVVSFVLMASSPRAADARYLMTRSDPLLSPIGPPYGKDPRFDRLYDIITKLLQNGGGDLEYQIKSQLDSGP

>RYamide/Luqin | Dpul11551

MARKESVFWLFCTLALMMSVVLVDAQTFFTNGRYGKRSEVRSRVASRSADERFFGGPRFGRSGNGGIVLGNSELDARNPERFFIGSRYGKRSEMEQIVPSPQVDESTSNSQEKETFLECNPIGIEQLYHCIERLKSAHHFDLMQHQQV

>SIFa | Dpul468

MRSSFIVVMVCVVVVLTFWGQVAEATRKLPFNGSIFGKRSNQGTDKLESPSNLQLLCDAAMNACSDWLPIGSK

>sNPF | Dpul3601

MELCPRINCWTTRTVLLVTFVVFLIHQDIQQNIASASPTPLLSGFEDYSEDRLNGEQPSLYELLLQREMLADKLDSEGRGHLIVRKSDRSPSLRLRFGRRADPDVPRVSAASNQHD

>Sulfakinin | Dpul16332

MPIIRQSSVKTLNLLLYIVRVVCAVDGQEEAQQQQQQQHRMKLTMLATVLAAVLVLGVGRATAAPADSSSTATGRRLLHSPNPTSHSKSIDSWLRWLLLRSRIGDKEKTKNGVPSNSFQLARSPVELGSSNPKLQAKLPPAIVQSNDDDETTGFGDEDFADEDVPLVLPEGRQAASKRQPDDYGHMRYGKRDFDDYGHMRFGRR

>Tachykinin | Dpul20413

MAVLMTVLAYCQPASAAATAVTDDDELMARQTRGLVLRSWRNAQQQTDHSADKTPSAKVAEPILPSQKEAMVFNGLPISMRLVLLQHLAGYDKRTPNSRAFLGMRGKKSSPPGADALTMEDNQLDDASGWPQGDILPDTYYFGPAPQKKKMHGEKFLGMRGKKMMNGLADGTAFIPNWRERYIYQEPFEKKRAPSSNSFMGMRGKRSESTTPTPNDYQFFNDDIIVDEELPDVDSKVSPRRSQERTPI

>TRH | Dpul9070

MRMEILQHHSACQRMLAVLLLLLSASSGFVPTADEQSPAVENWNDLVLRCRRSADDGSDSVTKEGSSLLPIRPVRRLGSEFLGKRAAAVVLTLENLCSELLFSEEEVDENWLCQCIRNWEQSTLPASSSLGGQHNMDEDSLAPAAAARVNKRGLGAILLSGKRMNRDKWNNNALTRRVMGSEFLGKRAIMGSEFLGKRAIMGSEFLGKRGYNGRSNGLSGPVKI

>Vasopressin/Oxytocin | Dpul9459

MAGLWTFCLIALSMTEMIIPLTAKPCFITNCPPGGKRSSQLVEPSSYLECAPCGPAGKGTCLGANLCCGSHFGCFFKTEETNVCLLTNLKSTQICNQHFWKTDLKSASCSLNGDKIDGICVADLLCCSLGNLPQDDL
