## Supplementary material for "Proteome-wide neuropeptide identification using NeuroPeptide-HMMer (NP-HMMer)": Figure S3_Tribolium_neuropeptidome.docx

**Figure S3**: *Tribolium castaneum* neuropeptides recovered using NP-HMMer.

>ACP | Tcas12692

MALKFRIFALVAVLVLMAWMFTGTQAQVTFSRDWNPGKRTENTDLHNTLKTASAVCHLLMNQVRQLASCDNNNELEPGATIFSGRR

>AKH | Tcas1325

MHRVLLTVLLITIVGLCAAQLNFTPNWGKRAPEGESNRCKESVDTIMLIYKIIQNEAQKLVDCEKFSN

>AKH | Tcas15624

MSRMFLIVVLIAFVGVCTAQLNFSTDWGKRSGSSAGSDANNCKEPVETIMLIYKIIQNEAQKIIECEKYA

>Agatoxin | Tcas15236

MKYTWLVLAACMVLVLAELLPGAAAGPYLDDDEGLPSDDDYTENAIDRLLQSAQKRSSLIYLFRRACVRRGGNCDHRPNDCCYNSSCRCNLWGSNCRCQRMGLFQKWG

>Allatostatin-B | Tcas12839

MRDAVAPVLGAVLLTCYSLQATLALSDETPLKSSNDNPQIEDEMSKRDWNKDLHIWGKRGWNNLHEGWGRKRSVPAWEEQQEKRAWQSLQSGWGKRFAPEDEYAIRQLAAMLDSQYDDYNPEIETNDDEKRNWGQFHGGWGKRSKWDNFRGSWGKREPAWSNLKGIWGKRSGEK

>Allatostatin-C | Tcas12020

MAAQLPRYLTKTLFIFLIATLVVANARPNHFGDASQVVGEPADGNNLLDSRLKPWELEMLVQRLSEISSQTGGDFAWDKSIRLPEAKRQSRYRQCYFNPISCFRK

>Allatostatin-CC | Tcas5980

MNRILMVVLESFLVAVLFEMKTDGFLIDRRSAASERNSDDYPDYQLGVKYDEYPMIVPKKRTALLVDRLMVALQQAIEEEEAANRVDGPPLTNSFQLSPEEVRKMDLQRRGHGSMSGQQKGRVYWRCYFNAVTCF

>Allatotropin | Tcas11663

MAFHHAALFFTLMFLWLLLANAQGRRDAKYPQVRTPQQRLTRGIEALKYHNMDLGTARGYGKRAVDMHNVNNFLLEWIALETRMRNLGIPRNLLRDQETIPE

>Bursicon alpha | Tcas5423

MKPLLSLVTLWKLTIFLSSMCLDPRLNSKIQVSGASTTDECQVTPVIHVLQYPGCVPKPIPSFACIGRCASYIQVSGSKIWQMERSCMCCQESGEREASVSLFCPKAKPGERKFIKVTTKAPLECMCRPCTGVEESAVIPQEIAGYADEGPLSNHFLKSHSQ

>Bursicon beta | Tcas6228

MFDKIILCLVYATCVYSVSEISEETCETLMSDINLIKEEFDELGRLQRICNGEVAVNKCEGSCKSQVQPSVITPTGFLKECYCCRESFLRERTITLTHCYDPDGVRLTAETVNSMDVKLREPAECKCYKCGDFSR

>Calcitonin | Tcas2952

MRPVLVLVMIFSVSYARYLEPYNYGYPRPLVDFLNRLSLSDKIKRCGNTFDESCANLPIIGASSDESWLAHSSPGKRCANVWGESCINGGIIGGGSDQSWLQGDDNPGRR

>CAPA | Tcas2233

MKTFLIYSACVVLFCIANCQGEPKEPKRNKLASVYALTPSLRVGRRSEGTDVKRRIGKMVSFPRIGRSESNWVPDDNSYGAQRPGANSGGMWFGPRLGRVQKRSENFTPWAYIILNGEAPIIREVHYSPRLGRESEEAYEEILDSNLDVL

>Pheromone biosynthesis activating neuropeptide (PBAN) | Tcas5741

MERFILINWTVLCVAVLFFETVLSTPHESSVPNERNDDSKETYFWFGPRLGRKKRNSSNDDLYQDMQKEELVSLTDALQDVPWAIIAVNDLLEGKRHVVNFTPRLGRESGEEFVNNAPEDRWLQNHETSGEMLYQRSPPFAPRLGRHSSPFSPRLGRENDRNLFS

>CCAP | Tcas12251

MTTAKLFVICIFAALAIETHSRFLPKSISKNLGATERVLEPKKRPFCNAFTGCGRKRSNLPALPEQSEVVDENLGSLLELSAEPAVEDLSRQIMSEAKLWEAIQEANMELHRRRQESAESSEEDAAVPARSATASCALPPCYI

>CCHa-1 | Tcas1150

MCHKQMTMSPLPIKLAKITVIVIFFCFAECAAGSCLSYGHACWGAHGKRNGHVPVREPSRDSGWFLSRLVQSPLYANEEAPSQQLFSDAQMEADPLKGQDDFRSMVDVYPNEENNFFDETFPNQRPRSHKNRVSKLLEKRSARMI

>CCHa-2 | Tcas10445

MNCWSTQVVLLAFVMAFVLAAAEAKRGCATFGHSCYGGMGKRTENNNEELLQDVQSEENPAFVFTGPRSENQQKLTPEQYDNISRVIRQWILSYRRAQEMRPDYN

>CNMa | Tcas12009

MRIAFGVIFVTGIFGGFFAENVFALPVARHHVITKDLDLDEMYKSYTSISNDRDDYNDKVTSRVYLDSPQISVSKNGKQQKTKQTAYLLVNTMSQRNKRYISYLTLCHFKICNMGRKRTSRYFHMIRQLDDNEA

>DH31 | Tcas11245

MVPNNGISLLMLLVAGMILFQATTTYAAPHSPSYPGYYSPISMEGQNPEYLLQTIARLRQALISDDDLENSKRGLDLGLGRGFSGSQAAKHLMGLAAANFAGGPGRRRRSEEEA

>DH44 | Tcas13396

MNKLSPESRSKRAGALGESGASLSIVNSLDVLRNRLLLEIARKKAKEGANRNRQILLSLGKRAFLQSRASGTYDNNV

>ETH | Tcas11466

MYLAIKNYVLKAAKNVPRIGRSNTNKNTNIDEMGKFFMKASKSVPRIGRRNENFDYGQPIVKRDEVPIWSDIADRFEYDPEILTSPEILEQLEMGDDPSVYEWEKIRTKRDSHKPHPKFYYVM

>Eclosion Hormone | Tcas1951

MLKLLFILLAVDTFFSNASIPVCITNCVQCKQMFGPYFQGRACGDACVSTNGRLVPDCNNAGTLGNFLKRLY

>Eclosion Hormone | Tcas8660

MDSGSRNFLVLLLLFASSLLVVDANPIGVCIRNCAQCKKMFGPYFEGQLCADACVKFKGKIIPDCEDITSIAPFLNKFE

>FMRFa | Tcas9228

MVPFAILILTLIIQLASGYNNEDFYSDNFDGFEEPSEEVSDMEVRRRNSNFLRFGRSGPNYEYEDYGEDFARPTRSGKIEKNDHFIRFGRSKQDFLRFGRNQPKATTNYLRFGRRNKRDTSNFLRFGRNSNFLRFGRNNESSYESPLVQLLSSLLKKEENKQRIV

>GPA2 | Tcas15086

MLACWLLFTLLSLSDAFMVKAVTARDAWQKPGCHKVGHTRKISIPECVEFHMTTNACRGFCESWAVPSGPKATPTQPVTSVGQCCNIMETEPVEARVLCVDGVRTLTFKSAVSCSCYHCKKD

>GPB5 | Tcas11033

MLGVQVWLFVGLSALVRCQSIIEAGLEPLDASGTIECHRRMYTYRVTQTDDNGKQCWDTLSVMACWGRCDSNEISDWRFPYKKSNHPVCVHYGRNRSVVTLRHCEEGANPSAARYEYLEAAGCKCQQCSSSDTSCEGLRYRPQRSHPASLGFRIN

>Insulin-like | Tcas1047

MNVPKLWLKVCFTLLLAGQIHANIDRKEFFCGKKLVKTLTELCAIYNYPTLPRRRFRRQIVDECCRSQCSRRYLVQYYCMEAHSSIAHLLKAKPEPEKPPERPVEAPKETPVGTTPSHISPNHGHCKCRKRRAKRINKSKRMQGQRNFIHNPVPPANIGHVERSQTPFYIWKFSRVY

>Insulin-like | Tcas4533

MSDTTRSENELELVFRDRSQSDWEEAWHKEKYTRCRETLIKHLYWACEKDIYRLTRRSDQSYNNYITNTDEEFPYLAPKKAKRLLRFRRGVNRRAGASITSECCKSSGCTWEEYAEYCPTNKRYTSYV

>Insulin-like | Tcas12331

MIVRKLLPATNKMDKRVLLFFFLINIIYVWSSPHMVHLMNKREIFCGTKLAETLAMLCKGNYYSPNPNPTKKSTNDIFAYNEYDEYFPNESDDENQLDFPFLQKEAVNSFLPIRFRRTRVGIVDECCRKPCSLKHLSLYCGQ

>Insulin-like | Tcas16130

MANFRGTKSKAVYCGRRLSETLSTVCKGNYNTLNKKSDIHEMGASRRPGYPSLSQHSLDYPYQSKANAASHHMSGFRRRKRRGVFNECCEKPCSLEELSQYCGGPSR

>ITP | Tcas12426

MNYRSSKSISTQAVWVCMVLAVVFQEITSSPAGRSPAFLPHHFTKRSFFDIQCKGVYDKSIFAKLDSICEDCYMLFREPQLHNLCRKNCFTTDYFKGCIDTLQRSDEEAQIQLWIKQIRGAELGGLGPSASPPNTS

>Myosupressin | Tcas8494

MQQYAFVAIVFGAVAVFLANASTTAYYVSCPPNDALEASTSLRHLCSFIEQAVNDNVVIQDEPFRRIVGRNVNPNTKRQDVDHVFLRFGRPFGL

>Neuroparsin | Tcas14866

MCPFHNFITIILVLTVTVIIFSDKGTAMIHLPCKRCATIQECNADPPQLCVFGENRDYCNRRVCSKGPGEKCGDRFNILGTCGEGLWCSNKDNRCHGCYIPTMACYPDD

>NUCB | Tcas359

MNRYIPFAFFLISVLQVCFAPPVTQNKDKDKEKGEDEIGGLEDYMEYHRYLQEVVNALESDPQFRQKLEKADETDIRSGKIAEELEFVSHHVRSRLDEIKRVELVRLKELTEKKRQLQQNIDLEDPSHHHLDHSNPHTFEIDDLKKLIAKTTADLAEADKKRREEFKQYELQKEFEKQEKLNHTNGEDREKLEREFREREEKHKKHEKLHEPGHKAQLEEVWKEQDQMQQEFDPKTFFMLHDIDGNGLWDQDEVKALFIKELQKMYAAGEPEDDMRERAEEMERMRESVFSEVDVNRDGFIDYEEFLAQTKRNDFQQDHGWQGLDERKPYTDEELEEYIRRHQAANQIPHGYPPQGYAQHPPPGYHPNVAVHPNGVPVQQYHPGQVPQQQFHPGQLSQQHPQQLQGHLPDLNTNEVYPQQRQQQYQQPPQHYQQQQYQQMNYQQHPQQVNYQQQHPQVPVQNQQQQHPQVPVQQNQQQHPQAVNQQQVPVQNQQQAANQQVPVQNQQIPVQNQQQQIPVQNQPQGVPQVQGNVQQGAPQASNPNLNNAQANQV

>Orcokinin-B | Tcas5177

MRFVTVAIVIFVAAAATEAAPKPAGLSGYSRRQWSRLFGRSVDPIDGDLIGRSVDPIDGDLIGRSVDPIDGDLIGRSLDRIGGGNLVGRSVDPIDGDLIGRSVDPIDGDDLIGRSLDRIGGGNLVGRSVDPIDGDLIGRSLDGIGGGNLVGRGVDPIDGDLIGRSLDIKRLLDGYRRKHNAYEEVIGQKTIRNFGVLQLGGGYGVAKRFAQPGKYDTKTEKHRRGPLNGLIPGGAFGRAARSCLKTNCLRLLKGQMDAKIHPYFSFDSDRQATKPGIYKTYQLLEME

>PDF | Tcas1987

MRCETFVAILVALGVVQSYPYDYKYLDRDYASPGAHQLASWIASQLRSKDYIQPQEGPILPYRLPVQGKRNSEVSNAIIGSEETQKMYRDGRK

>Phoenixin | Tcas2849

MTILRGWRYNVFIGGLVGVIALTLYPIAIEPMLNSEKYKKIQERNRRGIKQEDVQPGNMKVWSDPFGRK

>Proctolin | Tcas9933

MFDRKLVFALVFVVFATLAVEGRYLPTRSNGDRIEKLRELLKDLFENEVEKEEYQADAPPRWHPESKLFYKREAPAH

>PTTH | Tcas13749

MDIWKDKNYNFLDYDEVDDRCDNEICQNNFDDLIKRKVKDNEDMTYQSTTRLSPYYHPSRPMPCSCGIEFRVLDLGHQYYPRYLHSGVCKSELCGGPYRCIERHYKVRVLKQKDPRNPEIRPSMALPDTLKGTWLSETITVTVACECSV

>RYamide/Luqin | Tcas9661

MHARKLIVVLVYILTVLVSVAVSKRYTSEKRVQNLATFKTMMRYGRGGPSPNNKENKVNIRPRADAFFLGPRYGKRSGWSPNASLVYPVSTPLCGLDEDLSCAYTGISDLYRCTPRKGESEEFTTSSN

>SIFa | Tcas10871

MQLALAKVFSVCIVVIILTSWIEMTEATYRKPPFNGSIFGKRGATIEYDSASKALSAMCEIASEACQTWFPSQEK

>sNPF | Tcas6591

MQRYSAMKCLCAVTCIMIVVATVTSAAPSYADYDNNIRDLWEILLQKEAMDDKFAPGGPHQMVRKSGRSPSLRLRFGRRSDASMTPEAAFMMAQAVDHETN

>Sulfakinin | Tcas4071

MGMKSFFTGVFLISSVYLLFIHQFQNVSAAPGNANNVDSHRLRARPFARLTPRTQYSRIKAEPFNEFIVDDDDLFELSKRQTSDDYGHLRFGKRGEEPFDDYGHMRFGRSGSD

>Tachykinin | Tcas8509

MGNVCQTQHYYTITRVYLFSPLSILTMHSTTITTAVVLATIYVVCAAEDHHKRAPSGFTGVRGKKSIPDSAYSTGNSDSDSIPELKAVDIVSDLGAVDKRAPSGFMGMRGKKPFSLWEGTYPDGVFKRAPSGFMGMRGKKDMEFANYADEYIKRAPSGFMGMRGKKDYDSSSSQLDKRAPMGFMGMRGKKDYDEIADEKRAPSGFFGMRGKKMPRQAGFFGMRGKKYPYQFRGKFVGVRGKKASPDDYYNVDLDTVGQELDLNQLMLLLTENEGESDIWNGNNEVGQYSQK

>Trissin | Tcas16524

MNKNLLVVLIVIGVVWGEAQSCTSCGSECQSACGTRHFRTCCFNYIKKRSSDSLAVDPSLRLELWLAKSRNPYFQRNFLDSFLELPEGVNQDHDMTQ

>Vasopressin/Oxytocin | Tcas8222

MSTIITSIILLVLSESLVSGCLITNCPRGGKRSKFAISENAVKPCVSCGPGQSGQCFGPSICCGPFGCLVGTPETLRCQREGFFHEREPCIAGSAPCRKNTGRCAFDGICCSQDSCHADKSCASDDKSPIDLYTLINYQAELAGDK
