## Supplementary material for "Proteome-wide neuropeptide identification using NeuroPeptide-HMMer (NP-HMMer)": Figure S4_Example_output.docx

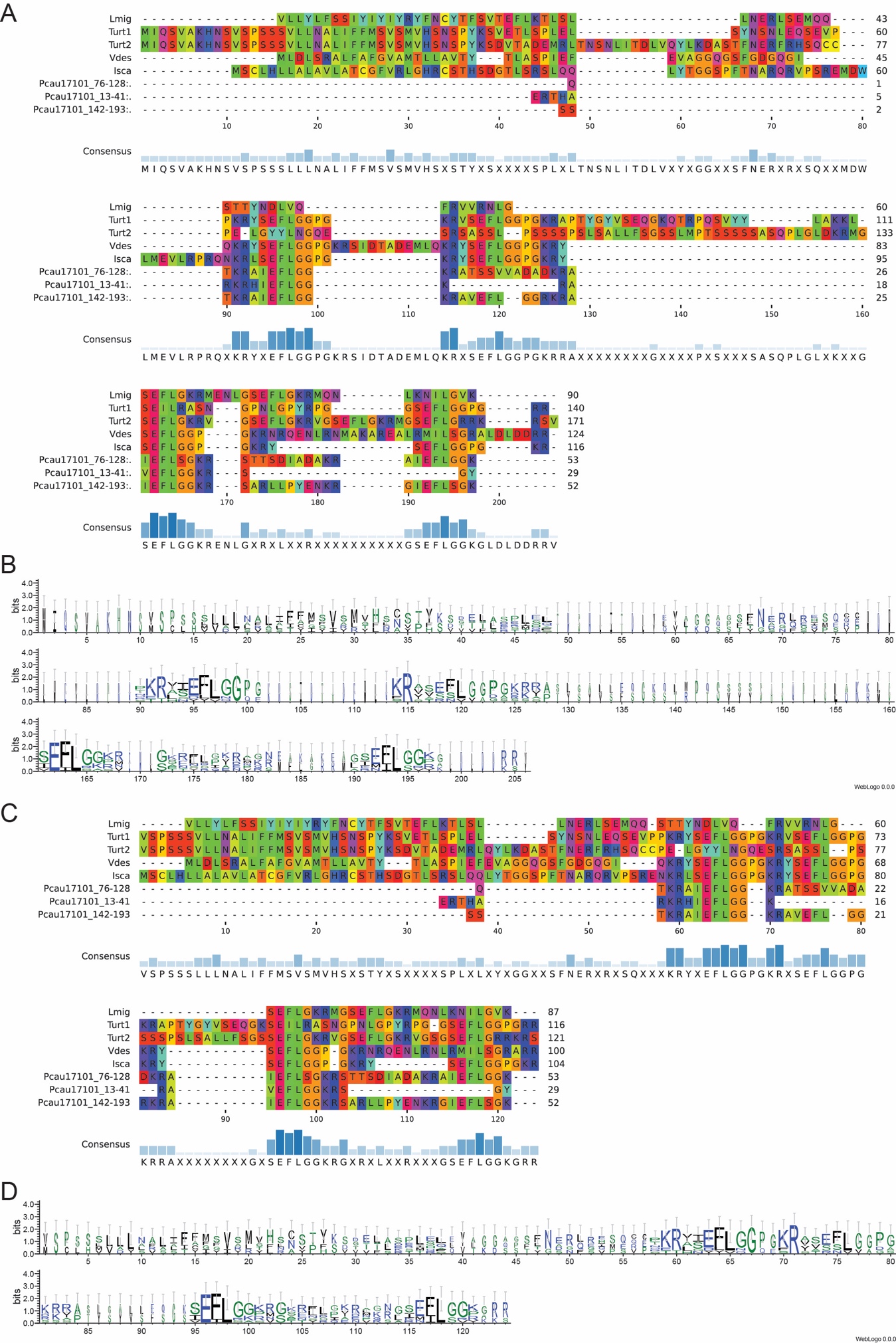

**Figure S4**: An example of a typical output from NP-HMMer. **(A)** non-trimmed sequence alignment and **(B)** corresponding sequence logo of thyrotropin-releasing hormone (TRH) in arthropods and a putative TRH identified in *Priapulis caudatus*. **(C)** A trimmed alignment and **(D)** corresponding sequence logo of the same sequences to better visualize precursors encoding multiple neuropeptides such as TRH.
