## Supplementary material for "Proteome-wide neuropeptide identification using NeuroPeptide-HMMer (NP-HMMer)": Figure S6_Priapulus_neuropeptidome.docx

**Figure S6:** Neuropeptidome of *Priapulis caudatus*. Highlighting indicates the following: predicted cleavage sites (red), signal peptide (blue), and predicted mature neuropeptides (green).

>Priapulus_caudatus_Agatoxin_1 | Pcau3479

MTLRFKVLALLFAIVLLGRAVSAQWLGDDDDDAEISDYLSDLYESDNDALQSLVKRTSRRMCLGHGRRCSLRRNDCCSRYICKCNLWGQNCKCEPRGFSFIG

>Priapulus_caudatus_Agatoxin_2 | Pcau17400

MQRIFAAFLALVLLVAVVASVPLDEEENALPDQEDLMALETYLQNIDKRSCMTRGRCSYSRNDCCRGYICKCNLFKQNCKCERKGIFGWGK

>Priapulus_caudatus_Allatostatin-B / Wamide | Pcau1995

MQMTCCHCYLLLLAGLAALALALAEDTNELGNAVDIDAPHDDKRGWHDLGHAYGKRAGWHDLQTSWGKRDNDADEQEKRGWHDLDTAFGKRGWRDLNTAFGKRGWRDLNTAFGKRGWRDLRMAFGKRDAVDTAEDQLVADEEKRGWHDLNTAFGKRGWRSLNSAFGKRGDNSAAIIEDLITEQLFDDIDSDGDKCIDRKELRLLVKRLLAGEL

>Priapulus_caudatus_Allatostatin-CCC_1 | Pcau1745

MASYPVSFTQLGVVLICFNMLLFACGRTLAYPTSALEQQLIDDPDKWSDVPETAERLSSPLRQYVLQELDRDSKGAGVGSSYRVQKRFGSVYQRPQRNRYSVRCLLNPITCFGRNQFRY

>Priapulus_caudatus_Allatostatin-CCC_2 | Pcau2514

MQQIHIFAGFFAFVLCISAVLGDHSTENELLSSPDSYRLLLDLIDNQIARAMAENSDMGHLAQKRDYFRKCQVNPISCFGGRQTRPDGRSPTSWWR

>Priapulus_caudatus_Bursicon alpha_1 | Pcau1435

MRCTLEGARRRLTYMKASTIVATLTLLLLTASSTTGDEVLPVEHQCQVRLFTQSTIGIFIMKLPGGVTCRNTQPVADFSECRGYCPSRDTFNTIVNDFESTCKCCASKVTRDVQIQLSCDNGTPTMHTFRQPVSCECKTCNLESKDAGWEVPGAVPEKKRK

>Priapulus_caudatus_Bursicon alpha_2 | Pcau5070 (partial)

MLVESGATVGSWRGNGVGSHWLGNDLCRLRRVIHVLRHAGCLPKSIPSFACQGRCSSHVQLRLDADLRLERSCMCCQEVGEREAHVRLKCAGDTVGYRRVKTYAPLECMCRPCA

>Priapulus_caudatus_Bursicon beta | Pcau15023

MQETRATVAWLLLLLLGLTVLDHAQSRAAVDRCHTLPSTLRMKKQVVNEEGKHIKTCVDTVAVSKCEGNCRSQESPSVMATSGFLKSCRCCREEQLSEKVVRLRNCYIPITGELIKSPLEVATHDVVIYEPTGCKCFRCLS

>Priapulus_caudatus_Calcitonin | Pcau17547

MLLSARPFPPATEIYDQHSAEIRRRSPLAPPLIASGLHHRQRQRHRHRHVSDHPDRVLGSLRRFLHACERQLLCRLMTSSTRRAMEQLALKRSRLLALQQLLKELDNSYETIQKRTCLMNGGMSHSCDYSEMSPGR

>Priapulus_caudatus_CAPA | Pcau12216

MREQLLEFVVRKAGADYARCVANGLKRSRSCKDPRPALRSYGPSLGMKKSSSLESLQAAVHDLARSEDVLDPGANYTRPQVKVVRGRGCNESFRAAVDRSYEAAIEAGVVLGPLSEEEQEAMRLRSDSVSGSLRSVPSVQAHAAERPHAVDDKRKKKKEKEKRKSLLRGLGSMFRVGRHKKDEEEAGNHAPNDVLADEAERQRQENLRCVPHPGRSDIGGRVTSHAGCVVCAAEPLPGVGGVEHGDVIPQLADEHETDGKAVGREASVDGDRRMSGDVEYGYVL

>Priapulus_caudatus_CCAP | Pcau11159

MLASAVRAATVLVCVVLLMVTRAAAEAEDAIYYDADMEPLRQLSNMRKRPFCNGFGGCRGAFGSKRGTDDVTDSADPDLANDVSGRLREAVIWRILGQKLRELQAMDHNEEREMEESAPAQQEERIFCRGFFDCMSRRRARRSTRQRFAVAEAK

>Priapulus_caudatus_CCHa | Pcau6983

MRTIEAVELCLIVTILACCLTATHGGCHAYGHACLGGMGKRADDAALQSSLQLSRDDTQRPRTAHGYYNQLRQLLQRVAALDRQKQTARMTSQNDDFRRDVGDYSDRFYPVDERRTYADDDGGIATGDLGKLIDQTRRLGGISKRWQQPLMDPSSMGDDSEDVEGINDVRWLRKRSISTSRRR

> Priapulus_caudatus_CNMa | Pcau438

MPSWLLLLVLSSAAVVCIVAIPSSNIDYESVPTRDQQTVDRGDAVIAYLLQALYDVYGRSDTQDAQSLVAKSYDVALHSPSAYRPPLSSSKRNMICYFKICKLGGRRRR

>Priapulus_caudatus_DH31 | Pcau15619

MGLALLLAVICIALPQVSAAAAGDRAKRDASIAEYIRALMEQLVQSEQENQQYKRAGIDMGFHRGYSGNQVAKHLLGLAAANYEGGPGKRR

>Priapulus_caudatus_DH44 | Pcau17368

MALLVLQLVFACLVISCATTSRQEARGYNDRHGSKTLLRLVPVLTPEKVDALEQARDADAVWRQAYEKRAGPRLSVISSLDVLRRQLQMVKDRQNKQAHGYQASANQNFLDRLG

>Priapulus_caudatus_ETH | Pcau13089

MRRATERNGVVVGWLLAFVVLLHTVSAQFYTKSGEGNLPRIGRRSDTFSLTDDAAPTFHSSYGAAQRHRFNDKLAGVIDRRGDIARVLTYLRRLENGNSDSF

>Priapulus_caudatus_Eclosion Hormone_1 | Pcau6991

MYLTTTKKVLRSPRCKIQSILMALLAAILMTSGQPITHQLAPSSVTSHADTKPFSQLTPTYGDTRELESIPREIEQKGNGNSPSSHNGWHSAALPQLDSVLLWRRYSRLVGDSALVGDKSAVISAPTAEQRSRTKPTHRHRKKRSAVICFFHCAECNKMYGAAAYNAPRCARHCTLSGGRSADVDCSNVTFWN

>Priapulus_caudatus_Eclosion Hormone_2 | Pcau7013

MNRAGGTAVLLLAFCTVGVYCHFPLMDCIEECFLCPVNVARGRDHMQMCANWCILTAGKSAGSGCQEMQHNWGINNENVIPPLELNALKNDISPSASREKRSNVLRSVLRGVRRRRQAAETKSETEREFAPLVPQQESTPKVTSNTVAGRSTHIVQRSRRADMYVVCLGQCVQCVLLYGYGTYDGKSCANACVLSGGASADPNCENSAFYIRFR

>Priapulus_caudatus_Eclosion Hormone_3 | Pcau4810

MQRTVEVLLVAAAVCVLTYPSYGVAGEIKESAPIEYIPMIDPVDTCLFKCESCYKGYGMLRCANECIETKGNISKKWRMRCGLFDDANLDILSLFNTIKRRK

>Priapulus_caudatus_GPA2_1 | Pcau14278

MGQESKAICFAIFAAVLCLPHSTSSSRNPWQKPGCHRVGFERTVTIPSCMPFTLATNGCRGYCTSFAVPSPQEVVAVNPTHAVTSYANCCNIGDSREVAVKVLCLDKVRTVTFKSAETCACSLCTKH

>Priapulus_caudatus_GPA2_2 | Pcau1435

MRCTLEGARRRLTYMKASTIVATLTLLLLTASSTTGDEVLPVEHQCQVRLFTQSTIGIFIMKLPGGVTCRNTQPVADFSECRGYCPSRDTFNTIVNDFESTCKCCASKVTRDVQIQLSCDNGTPTMHTFRQPVSCECKTCNLESKDAGWEVPGAVPEKKRK

>Priapulus_caudatus_GPB5 | Pcau14276

MLVTIYTLLGIASFTAHVSAKDTIDPTSTLICHKREYSYRISKPHNGLPCWDDVSVMSCWGRCDSNEIGDWVYPFKISHHPVCEHEVRIPRLVRLRHCHSLHPDPYYEVVDAESCACKACETKDTSCESIK

>Priapulus_caudatus_Insulin-like | Pcau4403

MRTDVNVATAVTSTTLMILIVFSGSTQGDSRLCGKRLTDTLMLVCMGRGFNWQVDVKRSAWPFMEKRGGSTDAGGQKRAHITPRGIVDECCKQSCSYDALESYCADRQGGPTATPGIAGLATLGRLSLSGMRRPSLSRTGLTRYTSLGRGIIALPSPMQPADNETSVQPEESTVNNESNLDNAHASSINTDPNRITAWKNQKFFFLPPPPS

>Priapulus_caudatus_Insulin-like | Pcau8412

MFQARTESEWRSVWHVETYRRCWHQLRPHISIACKKDIYRITKRTSPVEQNSTSSLASRIPLMIHKDGVFDINEVLTAKRAHSFLRSRGTPTRRVARNRNRDNIVTECCVQDRGCTWEEYAEYCPTHKRTRG

>Priapulus_caudatus_ITP | Pcau20214

MDTALRCCMWVGVALLMLITVTSALDPALREKLRMRHQNNIQESCGVISKLQLTLLDRVCEDCHMMFRIQNLHSLCRINCFQNYYFRSCAKALYQAPLLTYVERRQHGENSVLYD

>Priapulus_caudatus_Neuropeptide-F | Pcau20075

MSRTTILLSFAIGVVFVVIMCRSTMVAAGDMPLPPQRPGVFKTPHQLRTYLQKLNEYYAIVGRPRFGKRTFDDLREFVQRDSLEKPWYESILQGDRRHDY

>Priapulus_caudatus_NUCB | Pcau8615

MARLTWLILICLVASIVCPPVDKNSKKSEEDKKKENGTTDNNDIDLLEYDRYLKEIIGALEEDGDFKKKLESADFNDVQSGKIANELQFVNHNVRTKLDEIKRREMERLRQTTKKMLQKKEGRDVLKQGDIGGHIDHMNPRGFDMSDLAKLIQKTIADLETLDTQRKDNFKTYEMEKEYHHKEQVNNMTEEEKKIAEQEYLQQQEKHNDHPRVHHPGSKQQMQDVWTDNDHMDPGSFNPKTFFSMHDYNRDGYLDQKELEALFQTELDKLYDPDNPEDDMYERQEEMARMREHVQREVDTDSDGLISLAEFIAETHRSEWGEDEGWKDIRDEPVFDNEEYRAYEQFMAEQRHLASMDPQRLNTVPPHILEDFAKLPPDQQQQVMLQQHLQNQQQQQHPGQYQQHPGQVDPAQVDPRYQQHPGQVDQQYQQHHDQVDPRYQQHPDQVDQQ

>Priapulus_caudatus_Orcokinin | Pcau10184

MAKFLIVTCALVSIATRCRANQEHHEDHGMDKRLDSLSGFGLGEHKRQNLDLYSGSLLGGQKRLDSLSGFGLGSQKRLDSLSGFGLGHDKRSGDLDSLLGKTFGSDKSARELNKLALLGRMHKRFDPIEQAGFGSFAKRFDPIEQAGFGSFAKRFDPIEQAGFGSFAKRFDPIEQAGFGSFAKRFDPIEQAGFGSFAKRKREAIKSM

>Priapulus_caudatus_PTTH | Pcau124

MPCPDADVVLMLAAVLLVSVTVARPANKTIGADDDDVVVDKIECPPTPPDFLRTLLGPAFNERYMSVDEPADEWSADEWSADVGRRRDGKDGVLQDASHTTSAKPTHGPFVVGGDYERDLPTEKFYVDMFRSVLARMRDDARTPTTVATPVPDELEAVSDVGGNSTRGERTKRAAGATGGTPWGCDSRVVWHDLGPDRFPRFLRGVECTSSKCWFGHFSCRPKAFTVKVLRRKRDACSQAEVNGWRAPPGGDIFPTELLEKTWTFEERSVTFCCECVL

>Priapulus_caudatus_RYa/Luqin | Pcau12895

MVSLSFSSIVTFVLIALISVLIAADGTVAQWRPNTRYGKRSDDFPAGLTVYENDIVRRWDPQTRYGKRADFDTESKLNIAGVTNQDDNVSDESFSCVHTGVENLYRCFRKS

>Priapulus_caudatus_Tachykinin | Pcau19345

MESQWMIIAFLVSVMTTAGSEPTTDLYQEPQLQRRVEILDDIRHELNSLLQVLQQTTPRVTDMDDYTGQQDELQRFVTPADLIDQMTAESDKRAAFFGGRGKKYEMLADKRAAFFGGRGKKTGTMEDVEGELDDAKRAAFFGGRGKKSEMVMTEAEKRAAFFGGRGKKSDLESSEDELDAEKRAAFFGGRGKKSDAELNDEELEDAKRAAFFGGRGKKFDGGDDAKRSGFYGGRGKKLYPDDKRAGFFGGRGKKDDVDASETMVAIADDADEKRSGFYGGRGKKSGEEGKRGSGFYGGRGKRSHNLSVNM

>Priapulus_caudatus_TRH | Pcau17101

MWSDYNSLLPAEERTHARKRHIEFLGGKRAVEFLGGKRSGYHLLDNDDDVAQKRAIEFLAGKRSTAMSVADDGAAQTKRAIEFLGGKRATSSVVADADKRAIEFLSGKRSTTSDIADAKRAIEFLGGKRSAAAPADDDDDDSSTKRAIEFLGGKRAVEFLGGRKRAIEFLGGKRSARLLPYENKRGIEFLSGKRSASTTRLDDAIDAMLRTPSGRDERRRTLQLLDEIRRYDDAEAADDAAYGARF
