## Supplementary material for "Proteome-wide neuropeptide identification using NeuroPeptide-HMMer (NP-HMMer)": Figure S7_Adineta_neuropeptidome.docx

**Figure S7:** Neuropeptidome of *Adineta ricciae*. Highlighting indicates the following: predicted cleavage sites (red), signal peptide (blue), and predicted mature neuropeptides (green).

>Adineta_ricciae_Allatostatin-A | Aric33919

MQTQFLLIAGLLAILVLFHCNTVAESASLLHDDDPWDDISKEEFEIDKRLAAAKFASGLGRRLASEKFASGLGKRLASEKFASGLGKRLVAEKFASALGKRLAAEKFASGLGK

>Adineta_ricciae_Allatostatin-B / Wamide | Aric9711

MKVYFFVFVVLIVLVQTYFSNAEFSESSSENDNLNRFMNKRLSSQWGKRLQTIWGKRSASDNNELYQHILRELYRPSARMRQLEQSQNSVDNDPNHLTLEEFMAQR

>Adineta_ricciae_Allatostatin-CCC | Aric13105

MYQINLTFTYILQYPPDSLISDYRLALASDHLRQIERQQISPPSYILQSDYIEKLPHINWPKRDLRASLKTKRRMLCFFNTVTCFG

>Adineta_ricciae_Bursicon alpha | Aric10604

MVGLCLSLFLVLIFSSPIVALTSSTVQQCQVEIYNETLHFRHCLDHSISIQTTRCRGQCYSEEELVYDWEQTSTYRRYKRNFHCCVPKMTEAHETLVRCQNKQFQTVRYRLVTQCECKPCGDKCSEYF

>Adineta_ricciae_CAPA | Aric16547

MFISSLPSTFILASCCLAMIIFLKSSCAYVVDSNEIRQLDFNSPSYFDRYVRTTKFPRIGRSIYDNEDLSSLLDVNTNSEEATDENLEQRSVLFPRIGKRAFHNLLWANSRSNPHRMLDSQGRYYVNGYDYHIHQSPLNSVSRYRGKRSVSM

>Adineta_ricciae_CCHa_1 | Aric7312

MTRPSARKLPRCYSYFLVIVLLFQFIHSTQSLALSSLCGQFGHSCFGGNWGKREASTSTVKTDLLQANLDDLDGGTELTATANEDVLIKNLLLEEIRLSLLRQRLRHLLKLE

>Adineta_ricciae_CCHa_2 | Aric25886

MHLRSIFLAPFRLLIYLFIISSLFLLQTSRPIQAISLSRLCGSFGHSCFGANWGKRSSSDETSTRYIRLNLNDNDEARPVPAAGEDDILKEFRAKLLRQRLRQSASLDYDRIRRSLH

>Adineta_ricciae_FMRFa | Aric9742

MQSFSVYCLLLCSMIIFAAHQINAMETGDKFEATFPSEITSERTPEKVLTLREVLIHKARSAAAASSVSDDVAASAVAGGSDVEASVSVDGEEVDSVQCGNGQITIAKASLICLQLISTVTSDSNDASRSSQPNTLIQPSTIIDNEDIVNDDKEDDEDYFVVPRAAEFLRFGRQFPSTYSSFLRFGRSQPTFLRFGRPDPNGAAFLRFGRQAPSSASSNFLRFGRQFQQAASNNFLRFGREAQPAASSNFLRFGRKGEFLRFG

>Adineta_ricciae_Insulin-like | Aric22343

MLRSSFLMLCFLTILFISSTYEKNVTSRKRPHTSRHKPASIKLCGPTLVRMLDMVCDRARQLLMKTQRSSSDSTYQKRQMIVDDDPFTRTLSVTDYAQFNNTLVGDCCLQACTLKTLLKYC

>Adineta_ricciae_Neuropeptide-F_1 | Aric7400

MSFINNKFRCLISWLIFLSLAIQIAYAVSSFTDVPPPPKRPERFHSREELKRYLQEVHEYYAIIGRPRFGRALMKRYIDPLNTHLSYMFDNSDGNSKNPTNRMNTVVIPVMIHVVCKK

>Adineta_ricciae_Neuropeptide-F_2 | Aric9793

MLFTSSNTTRVSWIIFTVIAIQIAQVISAYTDVPPPPNRPERFHSREELKRYLQLVHEYYAIIGRPRFGRSLSSKYIDAQDRQLFDFFDVNGDNSIAPDEFYQRLENI

>Adineta_ricciae_Neuropeptide-F_3 | Aric30292

MSSINNKLRCLISWLIFLSLAIQIAYAVSSFTDVPPPPKRPERFHSREELKRYLQEVHEYYAIIGRPRFGRALMKRYIDPLDTHLLYIPNRSHEYSCHSGYDPCCFQKKSEHFFSLS

>Adineta_ricciae_Neuropeptide-F_4 | Aric14847

MNSFFTHFLSLLICSIILIRLTSSASFPDPPPIPPPNASPDDWIRFWKLLHSYYAIIARPRFGKRSELTMIATRPASRLELLFPRSPVSENEIIYTLSGNRYNANPNRRSSTGDIQTLIYSDRE

>Adineta_ricciae_Neuropeptide-F_5 | Aric24819

MNWFGVSIFVNLLLFSILSVHLTSSASFPNPPQKPPSNASPDEQALFWKLLHNYYAIIARPRFGKRSESFVKQPSYLETGHFDSAPSASSEYDMDRFNHPINVDSVYTFENSDKKRRRRR

>Adineta_ricciae_NUCB_1 | Aric11484

MKVTLTIISVFLLIFTIGAPPVLEKVNNDKAPVDTKGDESDEVLNNLEYGRYLKEVVEILENDPEFKRKIENASIDDIKSGNIAQHLSLVQHNVRTKLDEAKQREMIRLRELVAKKIRNLSEKERLELARAHPDAPHVKQFIPQHIDHQNVDGFNEKDLERLIQHASKDLDQLDRERERQFKEYEMQKEYERRHKLAQMTEEERKKAEALHEKAAEDKKHHPKVNHPGSVDQMEEVWENVDHLEANQFNPKAFFKLHDINGDGFLDEAEIEALMLKEAEKIHENNPEADPIEKQEEMDRMREHVMKEFDRDNDRMLSFNEFELGINGTGAKNDQGWQSIEDSTIYSEQEFNKFSENLDHAPTPSAPHQGAGASGNQIPPQDSAHLPRAQPPSINH

>Adineta_ricciae_NUCB_2 | Aric6212

MKLLIYLLHVCVLLAAVGAPPVVENARRGKAPAPSSNNEDSDDMIDKLEYGRYLKEVVEILESDPNFKRKLEKASADDIKSGNIAEHLSLVQHNIRSQLDEAKQREMARLRQLIGRKIRTLSDKQRNALARIDPNGEQIRKLLPQHLDHDNSDTFNEADLERLIRHASKDLDEIDRQREEEFKQYELRKEYQRRAKLAKLTKDERKRLEELHHEALEQKKRHRPVNHPGSRDQMEEVWTKVDKLETNQFKPKAFFKLHDINGDGFLDEGEIEAIMLQEAAKIHDGPADADPVEKEEEMERMRQHVMREFDKNNDRMLSFEEFEHGIDGKDAKNDQGWKSIEDNPVFSDQEYQRFAEHMPPISTSIPPQQTPSTSSSSVQAPVVSETVANASSVVPANQPLNIPRARPAAKTQ

>Adineta_ricciae_Orcokinin_1 | Aric20607

MTTATMSTVLIAFLIAALCQLGTMGDEEKKRTLDSLSGSDFFKKSLDSLSGNDFFKREDEKRTLDSLAGSDFFKKSLDSLSGNDFFKREEEKRTLDSLSGSDFFKKSLDSLSGNDFFKKSLDSLSGNDFFKKSLDSLNGNDFFKKSLDSLSGNDFFKRGVRPLGMLRYSGRHMHRHHSNDQN

>Adineta_ricciae_Orcokinin_2 | Aric22421

MTISINMSIMTVALMMITIYFVGINGEEEKKRTLDTVSGSDFFKKSLDSLSGNDFFKRNDEKRTLDSLSGSDFFKKSLDSLTGNDFFKREEKRTLDSVSGSDFLKKSLDSVSGNDFFKKSLDSLNGNDFFKKSLDSVSGNDFFKRRINDQQYHHPHMSRDHYHVRFHKSKPADDNDK

>Adineta_ricciae_PDF | Aric19641

MILFITLATFCPNQAESRAVHNIYKNGQPSHIAYLRLGDNDELASDDNSGENQDVYTHTKRKSELLSSLYGLPYALARKRQAN

>Adineta_ricciae_Proctolin | Aric30294

MHNIFSSIYLSIILSPIASNGARYHPRRAIDDDDDADDTRDLLRSLLRQASLRKIGFDSDDEDNYSNRFYFNEKIFFVQ

>Adineta_ricciae_Vasopressin/Oxytocin_1 | Aric7596

MHKFTVLILSVILVQLSYACYITNCPIGGKRSSSFKLNHLTHDCPRCGMNGQCFGPSICCSSLGCRIGHPSDIRQCSVESQSTVPCLIKGAVCSTVPNGRCAANRVCCGTELCRMDESCSIIANQDDE

>Adineta_ricciae_Vasopressin/Oxytocin_2 | Aric13180

MHQFAILILFVAILEFNHGCYITNCPIGGKRSSDLDNESYKHQCPRCGFGGQCFGPSICCNGLGCRIGHPSDVRQCSKESHSLTPCVINSAVCSSVPNGRCAAHGVCCGTESCQIDESCSEASNQEADNSREEHLPHSRFTLLQ
