## Supplementary material for "Proteome-wide neuropeptide identification using NeuroPeptide-HMMer (NP-HMMer)": Figure S8_Brachionus_neuropeptidome.docx

**Figure S8:** Neuropeptidome of *Brachionus plicatilis*. Highlighting indicates the following: predicted cleavage sites (red), signal peptide (blue), and predicted mature neuropeptides (green).

>Brachionus_plicatilis_ACP/APWGa | Bpli47440

MKILIGLSTICLSALILSIYIQESEASPQLTFSSDWSGGKRELEKSENENFDSIKSIEKSENENLDKRAPGWGKRAPGWGKRNFMDETKIDEIISRIKEYEHKATELRNFLFQMMEMHKNY

>Brachionus_plicatilis_AKH | Bpli5357

MNCRKVSNQALIITAVLLYFVGFSEQQFSYSANWGKRSNEENVENKCVSILSLKSYLLDVIPNLEQEIIVIQK

>Brachionus_plicatilis_Allatostatin-A | Bpli20339

MSILPNKVLILFSLMVLFSLFSNAKSENAEEKHEQSASEESEMEMEKRAIDKMRFYGGLGKRFDDNSYETMEYGDDEVELDKRLDRMRYFGALGKRSGELASNKLKTMKRLDKMKYFGGLGKRSDYYPMWRNSQLRYLLAKKSNFDRLRYLGNIGK

>Brachionus_plicatilis_FMRFa | Bpli3895

MNMNMNFYALIALLISSQIQAILSSDPQNDPESKKELEYMDKLRQLINIEKILYKMKKPELDENHLEADSYDELEPNADDYDFGNNEAKRSAYLRFGRSPAFLRFGRNQAYLRFGKSYGNFGAKRNPTYLRFGRSV

>Brachionus_plicatilis_Neuropeptide-F_1 | Bpli4070

MCAKNQVIISCIGLAVIAALIGQIEATEYPTPPPVPSQFLTPNDVQKYLNQLHNYYMVVGRPRFGKRSRVRAADYSDADDEPYIGSMFSFLDLNGDSAISLAEFKRIFDKESIAE

>Brachionus_plicatilis_Neuropeptide-F_2 | Bpli13495

MHVKIFLPILFGLILCQLIAETVSFGIRDTSDEMPERPAVFKSKKELIDYIKRVNEYYGKVGRPRFGKRNYPLFGSKFNQQYRDSNEWNFN

>Brachionus_plicatilis_Neuropeptide-F_3 | Bpli13883

MFAKSGFLKDFSNISKYSAAFILFLIVVSMCENVGAMDNQYSDLPPPPAKPERFTSKQQLKEYLVKLHEYYAIIGRPRFGRSQNLFNENIQSSELGTDAKENLIPVTLAINVMDFNGDGFLTKKELRNFVKLVEDYL

>Brachionus_plicatilis_NUCB | Bpli21098

MLKFVVLCSIIGLVVCPPVLPDNNHQPVHKSVNETLEKDPLLSLEYHKYLKEIVNILETDPEFKKQIENASIEDIKSGRIAEHLALVDPSIRTKLDEIKRMEIERLRRAIQRKAALSNLKAHEIQTLLPKHVDHGNIDTFEVHDLERLIKQATFDLEEVDKLRREEFKDHEIEKEYERRKKMQQADAETRQKLEEQHKRSLEMRKNHQRVNHPGSKDQLEEVWEEEDKLDGEDFEPVTFFKLHDIDGNNYWDEFEVEALFNIELDKIYNKTDPEFDPAERFEEMNRMREHVFGEIDKNKDRLVSLDEFVESTKSREFKQNEEWKGVDDEAQFSEDQIREYSQAHVEEERLMSTPVNDLRREQAHRQEDTQHHQPPRQHHQQPPEIQQHQPPQQHHQNVQHP

>Brachionus_plicatilis_Vasopressin/Oxytocin | Bpli49535

MNENFLMMNLNDKKNVKILFILVVINLNVINACYITNCPWGGKRSQPFLDFENAHQCRKCASGLGMCFGPRICCGPDMGCLIDTKETSVCQLEDIKSNVPCQPYGKICDKVEFGRCATSNLCCNPDHCLEDSTCVSEENDYSEDYKEIDTKLLKALKRLISKKRQKNENYESHNIDNYQS
