## Supplementary material for "Proteome-wide neuropeptide identification using NeuroPeptide-HMMer (NP-HMMer)": Figure S9_Didymodactylos_neuropeptidome.docx

**Figure S9:** Neuropeptidome of *Didymodactylos carnosus*. Highlighting indicates the following: predicted cleavage sites (red), signal peptide (blue), and predicted mature neuropeptides (green).

>Didymodactylos_carnosus_Allatostatin-A_1 | Dcar12455

MMMYNAIVIISIVVLFHINIVYNSTIHDARSNEETLQDHEHQISLIDQDDEDLYDMDKRLANMKFASGLGKRLANMKFASGLGKRLANMKFASGLGKRLANMKFASGLGKRLPSMKFASGLGKRNEYEDLMDFTDENENQFNRHYF

>Didymodactylos_carnosus_Allatostatin-A_2 | Dcar11442

MMYNAIIVISIVVLFHINVVYNTAIHDTKSDAETIQDHEHQTSLIEQDSDDDTYDMNKRLANMKFASGLGKRLANMKFASGLGKRLANMKFASGLGKRLPSMKFASGLGKRNEYEGLMNYIGENGNEFNRQYV

>Didymodactylos_carnosus_Allatostatin-B / Wamide | Dcar20486

MSLKLYNDQSVELIAKHQRIERRLSNVWGKRSLERRLSSTWGKRGISSNSDENVLDDLCEQYLLTRRLQEPASRKHEMQHLNSDESNPEVHELYRQPNYLEHTDQEPNEDTSIDETRNLSS

>Didymodactylos_carnosus_CCAP_1 | Dcar4055

MTSRWSLLSLLLFILFNIIQTSPVIVSDGNDPDSIDQLITTIDGNDNNIFGNDDANLSYQQKDTTHLIKNIMQETNPILQNLLLNQLREELNHMCVEGHFGPMITESCKRILNHLNQQPPLIRNKQQHNYQNKQQHNSDDVYSTDNTRQLKKRFFCNGFIGCKSGR

>Didymodactylos_carnosus_CCAP_2 | Dcar18136

MTSLWSLLSLLLFVIFNIVQITPVLMVDGSNDQDSLNPLLSVTDGIDAIDDVNSSNQQQNIGHIIKNIIQETDPIVQNLLLNQLREDLNRMCIQGHFGSTMAHACKHILSHTVQQQLSPLMREYQHKQQHNDDDIDSTTQTKQLQKRFFCNGFIGCKHGGR

>Didymodactylos_carnosus_FMRFa_1 | Dcar11992

MQPDRAPVPVDSEPDKNFIYELKSQHHDDDKSSSSENQDYLIDNINQRPFNSFISSNNYEDDEYQRSERAAAFLRFGRPSSSFLRFGRSNPSFLRFGRSNPSFLRFGRVPADKRTYQSSFLRFG

>Didymodactylos_carnosus_FMRFa_2 | Dcar19267

MTRSIIYYSEPNKNFVYDVKQHNDDKKINSENQDYLTDDSKQRPLNSFTSDNDNDNDDEYQRSERAAMQKFGPPSASFLRFGRSNPSFLRFGRSNPSFLRFGRSNPSFLRFGRTNLSPRLDWAPVEKRTQQASFLRFG

>Didymodactylos_carnosus_Leucokinin_1 | Dcar4543

MKIDMILPGVVCWIFLMLTVVHSFAFEHQIGKDVNADDEQKIHILEQHPSTLLSLLEDSHPTKFYPVYQESDINDFDDENDSSDDEHELEKRRFNAWAGKRMVPTKQRFNAWAGKRSIPFYLREGRRFNAWAGK

>Didymodactylos_carnosus_Leucokinin_2 | Dcar5599

MKIDMILPGVVCWIFLMLTFMQSSAVEHQDGKPINHSDEQEIHILRQDPSLLSKLENLHPTKFYSLFHENGHDGNQNSEELEFEKRRFNAWAGKRMVPIKRRFNAWAGKRSVPLHLREGRRFNAWAGK

>Didymodactylos_carnosus_Neuropeptide-F | Dcar32648

MYSLTSNLTLILFLMLFLIITVQSIQPFSDVPPPPPRPQRFYSRQQLKEYLQKVHEYYSIVGRPRFGRSDSSIRNKMKPKEAIAKSYTTLIDGIVDYLDMNKDGCITRNEYSERILSK

>Didymodactylos_carnosus_NUCB | Dcar4121

MIGIKFTLLCCLSIIFSVYAPPVNKSPSNNKGDSVNDEKEAKDILDDLEYARYLKEVVEILENDPKFKSMIDNATVDDIKSGNIAQHLSLVEHNVRTALDEAKQREIVRLRKLVAQKVRLLNDVHRRGGLRGTPNDHMKNIVPQHIDHHNLETFGVLDLEKLIRRASLDLDEIDKQREKEFKDYELRKEYDRREQLSHLSPEEQQKLEHQYQETLEKKKKHPKVNHPGSVDQMEEVWENVDHLEPEQFNPKTFFSLHDTNTDGFLDENEIEAIMLKEAEKIHNNATEVDAVEKQEELDRMREHVMREFDKNNDRMLSYQEFMSGINGTGAKNEQGWESIEDKPVYSDQEFQQFSDKMAHQSTPLPMHQQSQQHIERQQEHQAQQNPVPNNIPSPVNNQGYPGHQQQAQQNPVPNNIPLSVNNQGQQQQAQQNSAPNNIPSPVNNQGHPEQQHQAIHT

>Didymodactylos_carnosus_Orcokinin | Dcar5202

MESNESVVTLDVGELALFLFLLLFQKRTLDSLSGSDFFKKSLDSLSGNDFFKKSLDSLSGNDFFKREDEKRTLDSLSGSDFFKKSLDSLSGNDFFKKSLDSLSGNDFFKKSLDSLSGNDFFKKSLDSLSGNDFFKRSLDSLSGNDFFKRNDDLRTLYLIHELNLAHNGHRHPRHRFNKAQHHQED

>Didymodactylos_carnosus_Phoenixin | Dcar12179

MQDNLEGKKKLPRPDLRVPNPLLRTPETLTNLRSRILIIGTVLGLGIGIGVVFVYPYMNIEKFKRIQKTTRAELPPQETLQPAGLPVWRDPFDRKRTQG
