## Supplementary material for "Proteome-wide neuropeptide identification using NeuroPeptide-HMMer (NP-HMMer)": Figure S10_Rotaria_neuropeptidome.docx

**Figure S10:** Neuropeptidome of *Rotaria socialis*. Highlighting indicates the following: predicted cleavage sites (red), signal peptide (blue), and predicted mature neuropeptides (green).

>Rotaria_socialis_Allatostatin-A_1 | Rsoc7983

MQTKFILIVGLLAILFVFHCNTVESAVHLDSDEQLDDSSLEDSYELDKRLTSAKFASGLGKRLTSAKFASGLGKRLVSAKFASGLGKRLTSAKFASGLGKRLASAKFASGLGK

>Rotaria_socialis_Allatostatin-A_2 | Rsoc31746

MQTKFFIFAGLLAVLFLFHFNTVVESTSVANSDEQWNGISSEDYDFEKRLASAKFASGLGKRLSSAKFASGLGKRLSSAKFASGLGKRLASAKFASGLGK

>Rotaria_socialis_CAPA | Rsoc13646

MMFVSSSTITINSFYYILIILFIPSTCSYVLQSDDLKNSITDQQYGLNIPLYYDRFLRTTKFPRIGRSSSGTEEISSSDYQNMNVEENESSNNDNDSRWFEQRSVFFPRIGKRANNHVLWGNTLSSPYRMLDGHGRYHINGYDYHIRQNQPGSMTNYEGK

>Rotaria_socialis_CCAP | Rsoc11670

MSHHWLLLSIFLTIFLSQISNLHSISIDSGKSTYMIDENNGTQRAKDALIRLIENIPGESNPHKQKLLLNELREYLNRMCTVGYFGSLHAQACQRILNVIHEHDTNEENNDTTTNDQTRTEDHGIQKRFFCNGFIGCRNNAGR

>Rotaria_socialis_CCHa_1 | Rsoc3703

MNRQSIVLMPLCLSYFLIILLVLQSTRSVQSIALSSLCGDFGHSCFGGNWGKRELLRDRLVPDVVHTNTDTNGDGIEIPSTSNKNPMHNFLKKLRSQFLFQQRLRQLLQLE

>Rotaria_socialis_CCHa_2 | Rsoc22536

MNSPSFILDRIRFSIGLLLLTSLLFCQFPQPVQSISLSNLCGRFGHSCFGANWGKRTLPNELPQEYLLWNSNSNDGEATESAIMDRDPTNDYILEQIRLANEELFRHRLRQLLRLE

>Rotaria_socialis_FMRFa_1 | Rsoc12256

MYLYAIISSLICIQLLSIVSIDANPLNPALYSGAFIQRPTLLDSAATSNEDNQNEIYIQHPRLSSFLRFGRQMPSASASSLRFGRGGQIGGQVGGTFLRFGRQTQNPLSSNFLLSGNKGEFLRFG

>Rotaria_socialis_FMRFa_2 | Rsoc20036

MYLFTIISALICTELVIIASSESSEAIPTDQLNPLQPLKTFDNDFFEDSSQNDDENEYARRATSFLRFGRRGSPATGSFLRFGRDGHREGTFLRFGRSNPSFSRLGRSGNNGGKFLRFGREAQDESPSNNFRSGRKSDFLRFG

>Rotaria_socialis_GPA2 | Rsoc17844

MTVTLHLIFYAALLAAIMISRTYQCHLEFHTELLHLSGCLNTSIEIETTYCQGYCSSQDYLIYDWQSESRPYRHQHNITCCSPKMTVSREMKVLCDDRQPRIIKYPLVIRCQCRACTEICIS

>Rotaria_socialis_GPA2 | Rsoc26209

MVGIPIFLFLLIHINLSSTTTTLVKHHCQVEPYTETIYLQNCRSQPIKIDTTRCRGQCYSEDLLIYDWQNAPTHYRHKRHLFCCSPTNTEAQEIRVTCENENEPQTVKYRFVISCECKPCNDRCTE

>Rotaria_socialis_Insulin-like | Rsoc14244

MFRSSIFKFAIILIFMTSLINAKNTNNRRLKVLAIRQPTSIKMCGLALIHLLDTVCTRAEQILVRNRITTTLSSPTKRPRKTQDNLYSYATSIIDNTQFNDTLIYDCCLQVCTLKKLLKYC

>Rotaria_socialis_Leucokinin | Rsoc6591

MNTFVILTGILSCFLLMLSVTHAYTLTDKKNTADPQWLMDRPLAMARNKFFSMNQYDDDSMINSEEDDEHELEKRRFNAWAGKRSLVGKRRFNAWAGRR

>Rotaria_socialis_Neuropeptide-F_1 | Rsoc6251

MNSLVPLFIGLVAFFIISFQLASSANFPNPPSFPPPNASPEQWRAFWILLHNYYAIIARPRFGKRHDSVLSHLEHPSLVTTLNSNFASPSLYETDLNDIFGQNSRNANADQKAHFDELYTFVPTNRKQR

>Rotaria_socialis_Neuropeptide-F_2 | Rsoc17324

MNSISLSLLVNLLILLVVSFHLTSSASFPNPPQRPPSNASPDEQALFWKLLHNYYAIIARPRFGKRFSPASSYSKHAPFLPTANPDSIGLLSSAYNTNRMLNKNIPNANDQRTDDDNFYVLEYADQRRRRRL

>Rotaria_socialis_Neuropeptide-F_3 | Rsoc34079

MPNHKDIITKHVSWIVFIFLAIQTAQVLSAFTDVPPPPKRPERFHSREELKRYLQLVHEYYAIIGRPRFGRSTQLKPRIESSDFDLFKFFDTNGDKSISYKEFHGRIGA

>Rotaria_socialis_NUCB_1 | Rsoc4608

MKLKISIFFFFAVIIIIGAPPVVEHPAEDKAQTTPQDPQGSDDVLDNLEYGRYLKEVVEILESDPEFKKKIENASLDDIKSGNIAEHLSLVQHHIRTQLDEAKQREMIRLRELVGQRARNLSDKQRAALARSDAGGKLTRELLPQHIDHKNVETFGQTDLERLIRHASRDLDELDRKREKEFKEYEMQKEYERRAELAKLSPEERQKIEATHAEALEKKKHHPKVNHPGSVDQIEKVWEDVDKLDADQFSPKSFFNLHDINTDGFLDEAEIEAIMLKEAEKVHEGTPEADPVEKQEELDRMRQHVLTEFDKNADRMLSLEEFLVGINGTGAKNDQGWQSIEDSTVFSDQDFNKFSEKMAPVSTSIPIHQTPSLPNNQQVPIHPEAAVERPAAGQQQIHIPRAPPAVNNP

>Rotaria_socialis_NUCB_2 | Rsoc36363

MKLAISILSIFFLITTIGAPPVGENVNKNKKTVDSKEDSSDEALNNLEYSRYLKEVVEILENDPEFKKKLENASLDDIKSGNIAQHLGLVQHHVRTKLDEAKQREMDRLRELVGRRIRNLSEKERMELARGNPNGKNIKDLLPQHIDHQNSENFAEADLERLIRHASKDLDEIDRQREREFKEYEMRKEFERRAKLTEMNEEERKKQETLHQEAVEKKKHHPKVNHPGSVDQMEEVWEDVDHLEADQFNPKSFFKLHDVNSDGFLDEGEIEAIMLKEAEKIHENTPEADPIEKQEEMDRMREHVMKEFDKNNDRMLSFDEFEHGMNGTEAKNDQGWQSLEDSTPYSDQEFQSFSDQLIHARGSDASQPHNPPPANAVPSSPQEHIPRAPPSMNSH

>Rotaria_socialis_Orcokinin_1 | Rsoc26551

MTTTNMSMMIVALVITTMCLLIANGEEGKKRTLDSLSGSDFFKKSLDSLSGNDFFKREDEKRTLDSLSGSDFFKKSLDSLSGNDFFKRDDEKRTLDSLSGSDFFKKSLDSLSGNDFFKKSLDSLSGNDFFKRSTKAREYAIAELLRHSLYPHMTREHFRTRFNKQKN

>Rotaria_socialis_Orcokinin_2 | Rsoc29739

MTAATMSTIYVALVIATICQLGTMGEEDKKRTLDSLSGSDFFKKSLDSLSGNDFFKREDEKRTLDSLSGSDFFKKSLDSLSGNDFFKREEEKRTLDSLSGSDFFKKSLDSLSGNDFFKRNLDSLGGNDFFKKSLDSLSGNDFFKRNMKPSEFNMLSERRSHHPYRVHERFSVINNQDK

>Rotaria_socialis_Phoenixin | Rsoc23864

MNQQQTNKRLPRPDLRISPSQWSAAQKGGTSSRVAIVATVTAVIGGLSLVFLYPYMNIEQFRSVQKANRAGIHAEDVQPTGLPIWRDPFDRKKSA

>Rotaria_socialis_Vasopressin/Oxytocin_1 | Rsoc23386

MDQLIFIVLSVAIIQLSCACYITNCPIGGKRSLLLGNNLISHQCPRCGLNGQCFGSSICCTGLACRIGHPFDVRQCSLENRSVTPCDVNTSICSAVSNGRCAANGVCCSADSCQMDKSCLVSNQERDDSSREDYLAQAEIMLFE

>Rotaria_socialis_Vasopressin/Oxytocin_2 | Rsoc39311

MRQSTLFILSLAIIQLSYACYITNCPIGGKRSLFASSILHDHQCPRCGANGQCYGSSICCTSSGCRIGHRSDIRQCSIEDHSVIPCTIKSASCSVLPNGQCAANGVCCNTESCQMDETCSMSSNQNDDSLQEQSHVL
