## Supplementary material for "Proteome-wide neuropeptide identification using NeuroPeptide-HMMer (NP-HMMer)": Figure S11_Protein alignments.docx

**Figure S11:** Multiple sequence alignments of protein hormones discovered in Priapulida and Rotifera. Alignments of **(A)** burscion alpha, **(B)** bursicon beta, **(C)** glycoprotein hormone alpha 2 (GPA2), **(D)** glycoprotein hormone beta 5 (GPB5), **(E)** insulin-like and **(F)** prothoracicotropic hormone (PTTH) neuropeptides. Complete alignment for only the insulin-like peptides is shown and the alignments have been truncated for other proteins. Arthropod species are colored in blue, priapulids in black and rotifers in red. Species names: Pcau, *Priapulis caudatus*; Rsoc, *Rotaria socialis*; Aric, *Adineta ricciae*; Dpul, *Daphnia pulex*; Bmor, *Bombyx mori*; Tcas, *Tribolium castaneum*; Dmel, *Drosophila melanogaster*; Isca, *Ixodes scapularis*.

(A) **Bursicon alpha**

**Pcau -----------DEVLPVEHQCQVRLFTQSTIGIFIMKLPGGVTCRNTQPVADFSECRGYCPSRDTF------NTIVNDF
Aric ------------LTSSTVQQCQVEIYN---------ETLHFRHCLDHSISIQTTRCRGQCYSEEELVYDWEQTSTYRRY
Dpul ------------------DECQLTPVI---------HVLQYPGCIPKPI--PSFACTGKCTSYVQV-----SGSKLWQT
Dmel -QPDSSVAATDNDITHLGDDCQVTPVI---------HVLQYPGCVPKPI--PSFACVGRCASYIQV-----SGSKIWQM
Bmor ----------HEVQLPPGQECQMTAVI---------HVLKHRGCKPKAI--PSFACIGKCTSYVQV-----SGSKIWQM
Tcas LDPRLNS-KIQVSGASTTDECQVTPVI---------HVLQYPGCVPKPI--PSFACIGRCASYIQV-----SGSKIWQM

Pcau ESTCKCCASKVTRDVQIQLSCDNGTPTMH-----TFRQPVSCECKTCNLESKD-------AGWEVPGAVPE--------
Aric KRNFHCCVPKMTEAHETLVRC-----QNKQFQTVRYRLVTQCECKPCGDKCSEYF------------------------
Dpul ERSCMCCQESGEREATVSLLCPKAAPGEPKLRRVVTRAPVDCMCRPCTALEESAVMPQEIARFLDDGSFPFKL------
Dmel ERSCMCCQESGEREAAVSLFCPKVKPGERKFKKVLTKAPLECMCRPCTSIEESGIIPQEIAGYSDEGPLNNHFRRIALQ
Bmor ERTCNCCQESGEREATVVLFCPDAQNEEKRFRKVSTKAPLQCMCRPCGSIEESSIIPQEVAGYSEEGPLYNHFRKSL--
Tcas ERSCMCCQESGEREASVSLFCPKAKPGERKFIKVTTKAPLECMCRPCTGVEESAVIPQEIAGYADEGPLSNHFLKSHSQ**

(B) **Bursicon beta**

**Pcau ----SRAAVDRCHTLPSTLRMKKQVVNEEGKHIKTCVDTVAVSKCEGNCRSQESPSVMATSG
Dpul --SKTNLMSGTCETLPSTIHITKEEYTDGGILSRTCEGDIGVAKCEGSCSSQVQPSVVHPSG
Dmel LRYSQGTGDENCETLKSEIHLIKEEFDELGRMQRTCNADVIVNKCEGLCNSQVQPSVITPTG
Bmor --------EENCETVASEVHVTKEEYDEMGRLLRSCSGEVSVNKCEGMCNSQVHPSISSPTG
Tcas ---VSEISEETCETLMSDINLIKEEFDELGRLQRICNGEVAVNKCEGSCKSQVQPSVITPTG

Pcau FLKSCRCCREEQLSEKVVRLRNCYIPITGELIKSPLEVATHDVVIYEPTGCKCFRCLS---
Dpul FLKECMCCRESFLRERVVTLTHCYDANGNRLTG--K-SSSLDVKMREPADCKCFRCGDSAE
Dmel FLKECYCCRESFLKEKVITLTHCYDPDGTRLTS--PEMGSMDIRLREPTECKCFKCGDFTR
Bmor FQKECFCCREKFLRERLVTLTHCYDPDGIRFED--EENALMEVRLREPDECECYKCGDFSR
Tcas FLKECYCCRESFLRERTITLTHCYDPDGVRLTA—ETVNSMDVKLREPAECKCYKCGDFSR**

(C) **Glycoprotein hormone alpha 2 (GPA2)**

```
Pcau_1 ------------------NPWQKPGCHRVGFERTVTIPSCMP--FTLATNGCRGYCTSFAVPSPQ
Rsoc_1 -------------------------CHLEFHTELLHLSGCLNTSIEIETTYCQGYCSSQDYLIYD
Rsoc_2 -----------------TTTLVKHHCQVEPYTETIYLQNCRSQPIKIDTTRCRGQCYSEDLLIYD
Dpul   EPRNQPKGSISVSTRSSTSRGELSGCHQVGHTRRVTIPDCVS--FMITTNACRGFCESWSVPSSW
Dmel   -------------NSMGKDAWLRPGCHKVGNTRKITIPDCVE--FTITTNACRGFCESFSVPSIP
Bmor   --------------------WRKPGCHRIGHTRNISIPDCVE--FKITTNACRGYCESWSLPSIM
Tcas   ---------FMVKAVTARDAWQKPGCHKVGHTRKISIPECVE--FHMTTNACRGFCESWAVPSGP

Pcau_1 EV-----VAVNPTHAVTSYANCCNIGDSREVAVKVLCLD--KVRTVTFKSAETCACSLCTKH---
Rsoc_1 WQSES------RPYRHQHNITCCSPKMTVSREMKVLCDD-RQPRIIKYPLVIRCQCRACTEICIS
Rsoc_2 WQNAP------THYRHKRHLFCCSPTNTEAQEIRVTCENENEPQTVKYRFVISCECKPCNDRCTE
Dpul   EA-----LLKNPEKVITSVGQCCNIMASEDVTVRVMCLG--GPRDFTFKSAKTCSCFTCKKD---
Dmel   MMGSSLSVLFKPPKPVVSVGQCCNMMKSEEIQRRVLCIE--GIRNVTFNSALSCSCYHCKKD---
Bmor   LG--------FKRHPVTSLGQCCNIMEAEDVPVKVLCLD--GERNLIFKSAVSCACYHCQKE---
Tcas   KA--------TPTQPVTSVGQCCNIMETEPVEARVLCVD--GVRTLTFKSAVSCSCYHCKKD---
```

(D) **Glycoprotein hormone beta 5 (GPB5)**

**Pcau -----------DTIDPTSTLICHKREYSYRISK-PHNGLPCWDDVSVMSCWGRCDSNEIGDWVYPFKISHHPV
Dpul --AMFEQPDSQSKEQMDATPTCFRRPYTFKVYQEDSEGRSCWDVVTVTSCWGRCSSNEIADWRFPFKRSQHPV
Dmel --SSLSEIKPMNNGHIVTPLGCHRRVYTYKVTQSDLQGHECWDYVSVWSCWGRCDSSEISDWKFPYKRSFHPV
Bmor ----------------SMSVRCKLKRHSHKVMQTDLNSRRCWDDVKIVSCWGYCLSYEISDWQFPYKESHHPV
Tcas QSIIEAGLEPL---DASGTIECHRRMYTYRVTQTDDNGKQCWDTLSVMACWGRCDSNEISDWRFPYKKSNHPV

Pcau CEHEVRIPRLVRLRHCHSLHP--DPYYEVVDAESCACKACETKDTSCESIK----------------------
Dpul CQHDSILPRAITLRNCDPEVNLGTELYMALDAVTCRCELCRTDTTNCEGPHYDRRITTRRLN-----------
Dmel CVHAQRQLVVAILKNCHPKAEDSVSKYQYMEAVNCHCQTCSTQDTSCEAPANNEMAGGSRAIMVGADTKNLDY
Bmor CVHGERRHASVKLRNCDPGVEPGTEIYHYVEAVNCRCQVCSSEDTSCEWLPPDSSLLGGLILKEELE-EELE-
Tcas CVHYGRNRSVVTLRHCEEGANPSAARYEYLEAAGCKCQQCSSSDTSCEGLRYRPQRSHPASLGFRIN------**

(E) **Insulin-like**

```
Pcau   MRTD--VNVATAVTS------TTLMIL---------------IVFSGSTQGDSRLCGKRLTDTLMLVCMGRGF--NWQVDVKR-----------------------
Aric   ------------MLRSSFLMLCFLTILFISSTYEKNVTSRKRPHTSRHKPASIKLCGPTLVRMLDMVCDRARQLLMKTQRSSSDSTYQKR-------Q------MI
Rsoc   ------------MFRSSIFKFAIILIFMTSLINAKNTNNRRLKVLAIRQPTSIKMCGLALIHLLDTVCTRAEQILVRNRITTTLSSPTKR-------P------RK
Dpul_3 MA--HLTVG-------RCWMACIILL--AL---ATFTL---ARPPQENQPMTIRFCGRDLIRAIDEVCVAIKSPAAF-IDPVALNQS------SSL------ADSE
Dpul_4 MR--TLTAG-------KRLKTSVLLM--VL---AGLGLTQGRPPHQQNQPMMMRFCGIELTLVIDEVCTAHKSPA-F-VDHFILNRP------SSN------NEST
Dmel_1 MFSQHNGAAVHGLRLQSLLIAAMLTA-------AMAMVTPTGSGHQLLPPGNHKLCGPALSDAMDVVCPHGFNTLPRKRESLLGN-----SDDDEDTE----QEVQ
Dmel_2 ------------MSK-PLSFISMVAV--IL------------LASSTVKLAQGTLCSEKLNEVLSMVCEE-YNPVIPHKRAMPGAD----SDLDALNPLQFVQEFE
Bmor_2 ML------VSNAIQL------TVFMVAVLW--KNEA---------EAATKASVKFCGRHLSEIMSRVCHA-YNGPAW-----------------------------
Tcas_1 MIVRKLLPATNKMDKRVLLFFFLINIIYVW--------SSPHMV--HLMNKREIFCGTKLAETLAMLCKGNYYSPNPNPTKKSTNDIFAYNEYDEYFP----NES-
Tcas_2 ------------MDLQ----YVLVVVATVL--AGIHTCRTDEMANFRGTKSKAVYCGRRLSETLSTVCKGNYNTLNKKSDIHEMGA----SR-RPGYP------S-

Pcau   ----S----------AWPFMEKRGGST--DAGGQKRAHITPRGIVDECCKQSCSYDALESYCADRQGGPTATPGIAGLATLGRLSLSGMRRPSLSRTGLTRYTSLG
Aric   ---------VDDD--PFTRTL-----------SVTDYAQFNNTLVGDCCLQACTLKTLLKYC--------------------------------------------
Rsoc   ---------TQDN--LYSYAT-----------SIIDNTQFNDTLIYDCCLQVCTLKKLLKYC--------------------------------------------
Dpul_3 --------SADTQLRK----------------WTKLEDEKSDSLLRQCCVIGCTDNDLTTFCQIDRNQIMG------AAMHLRSQEWDMD---WLYEFLPRYGDVP
Dpul_4 -------------GNG----------------RTELDDKKSDSLLRQCCVIGCTEDDLATFCRIDRNQIRE------VAIRLRSEDVDAD---WLYEFLPRYGAIH
Dmel_1 -DDSSMWQTLDGAGYSFSPLLTNLYGSEVLIKMRRHRRHLTGGVYDECCVKTCSYLELAIYCLPK-----------------------------------------
Dmel_2 EEDNSISEPLRSALFPGSYLG-GVLNS---LAEVRRRTRQRQGIVERCCKKSCDMKALREYCSVVRN---------------------------------------
Bmor_2 ---------------DVPTVEQ---------PGGLLRRKRQLGIADECCLMGCTWEQLSEYCSIIAYSESPLEDLESHVIADRSAE----QE-----------NLA
Tcas_1 ---------DDENQLDFPFLQKEAVNS---FLPIR-FRRTRVGIVDECCRKPCSLKHLSLYCGQ------------------------------------------
Tcas_2 ---------LSQHSLDYPYQSKANAAS---HHMSGFRRRKRRGVFNECCEKPCSLEELSQYCGGPSR---------------------------------------

Pcau   RGIIALPSPMQPADNETSVQPEESTVNNESNLDNAHA---------------------------------SSINTDPNRITAWKNQKFFFLPPPPS---------
Aric   ---------------------------------------------------------------------------------------------------------
Rsoc   ---------------------------------------------------------------------------------------------------------
Dpul_3 TNA------------------------------------------------------------------------------------------------------
Dpul_4 APE------------------------------------------------------------------------------------------------------
Dmel_1 ---------------------------------------------------------------------------------------------------------
Dmel_2 ---------------------------------------------------------------------------------------------------------
Bmor_2 AGAKTTTTTTTPAVV---GSDEHVHVRGETGLASSHGYGRARGRRCWCRRKRRSGRRRASLAIIKNAVRAAPVVGTVSPLITWGRTLNTDLPRPDNDRYAYVVYT
Tcas_1 ---------------------------------------------------------------------------------------------------------
Tcas_2 ---------------------------------------------------------------------------------------------------------

```

(F) **Prothoracicotropic hormone (PTTH)**

**Pcau RPANKTIGADDDDVVVDKIECPPTPPDFLRTLLGPAFNERYMSVDEPADEWSADEWSADVGRRRDGKDGVLQDASHT---TSAKPT---------HGPFVVGGDYERD------------LPTE-------KFYVDMFRSVLARMRDDARTPT
Dmel -----------------------------------------------------SQYASDE-----GLDEMVGLRSLEHRAEEQQPDRSTSKMLSALFGFSPSTP--HPTE------MAMVMPH----QLPPMYYNDFYEDLV-----------
Bmor ----------------------------------------------------------------------FVPKAVA---LKRKPD---------VGGFMVEDQRTHKSHNYMMKRARNDVLGDKENVRPNPYYTEPFDPDTSPEELSAL-IV
Tcas ----------------------------------------------------------------------------------------------------------------------MDIWKDK------NYNFLDYDEVDDRCDNEIC-QN
Isca ---------------SPKPACSRATREDLERLLGSAFNARYMAIDKPDPEPTSSRLTARHSE-----D------PLS---DQADLE---------AGGFSVDADFRQD------------LPGE-------RRR-------------------

Pcau TVATPV------PDELEAVSDVGGNSTRGERTKRAAGATGGTPWGCDSRVVWHDLGPDRFPRFLRGVECTSS-----KCWFGHFSCRPKAFTVKVLRRKRDACSQAEVNGWRAPPGGDIFPTELLEKTWTFEERSVTFCCECVL---------
Dmel --------------------------------TTKRNDVHSAGCDCKVTNELVDLGGLHFPRFLMNAVCESGAGRDLAKCSHGSNCRPLEYKVKVLAQTSQSDHP---YSW-----------MNKDQPWQFKTVTVTAGCFCTK---------
Bmor DYANMIRNDVILLD---NSVETRT--RKRGNIQVENQAIPDPPCTCKYKKEIEDLGENSVPRFIETRNCNKTQQ---PTCRPPYICKESLYSITILKRRETK---SQ--------ESLEIP-NELKYRWVAESHPVSVACLCTRDYQLRYNNN
Tcas NFDDLIKRKVKDNEDMTYQSDVIG--KKTTRLSPYYHPSRPMPCSCGIEFRVLDLGHQYYPRYLHSGVCKS------ELCGGPYRCIERHYKVRVLKQKDPRNPEIR--------PSMALP-DTLKGTWLSETITVTVACECSV---------
Isca --------------------------------VREKRSETAKPWGCTSRLEWEDLGDDRFPRYLRNVKCLGG-----DCWFGKFRCKARAFTVKVLRRKSGKEKDGDCV---VSASTADLP-AELREHWEFEERAVAFCCDCSVDD-------**
