## Supplementary material for "Proteome-wide neuropeptide identification using NeuroPeptide-HMMer (NP-HMMer)": Figure_S5.docx

Adipokinetic hormone (AKH), Adipokinetic hormone/corazonin-related peptide (ACP), and Corazonin (CRZ)

```
Drome_AKH  QLTF--SPDW--G
Bommo_AKH1 QLTF--TSSWG-G
Bommo_AKH2 QLTF--TPGWGQG
Trica_AKH1 QLNF--STDW--G
Trica_AKH2 QLNF--TPNW--G
Rhopr_AKH  QLTF--STDW--G
Dappu_AKH  QVNF--STSW--G
Bommo_ACP  QITF--SRDWSGG
Trica_ACP  QVTF--SRDWNPG
Rhopr_ACP  QVTF--SRDWNAG
Nasvi_ACP  QVTF--SKGWGPG
Anoga_ACP  QVTF--SRDWNAG
Drome_CRZ  Q-TFQYSRGWTNG
Bommo_CRZ  Q-TFQYSRGWTNG
Rhopr_CRZ  Q-TFQYSRGWTNG
Dappu_CRZ  Q-TFQYSRGWTNG
Rhimi_CRZ  Q-TFQYSRGWTNG
```

Allatostatin C (Ast-C), Allatostatin CC (Ast-CC), and Allatostatin CCC (Ast-CCC)

```
Drome_AstC   ----------QVRYRQCYFNPISCF-
Bommo_AstC   ----------QVRFRQCYFNPISCF-
Trica_AstC   ----------QSRYRQCYFNPISCF-
Glomo_AstC   ----------EAKYKQCYFNPISCF-
Anoga_AstC   ----------QIRYRQCYFNPISCF-
Dappu_AstC   --------SKQLRYHHCYFNPISCF-
Drome_AstCC  --IQPSGSGGGRAYWRCYFNAVSCF-
Bommo_AstCC  ----GQSNNNRGRVLRCFFNAVTCF-
Trica_AstCC  GHGSMSGQQKGRVYWRCYFNAVTCF-
Rhopr_AstCC  -----GQQKGGRIYWRCYFNAVTCF-
Nasvi_AstCC  ------GQAKGRVYWRCYFNAVTCF-
Dappu_AstCC  ------GQSSQRVFWRCYFNAVSCF-
Rhimi_AstCC  ------------MFWRCYFNAVSCF-
Rhopr_AstCCC -----------SYWKQCAFNAVSCFG
Locmi_AstCCC -----------SYWKQCAFNAVSCFG
Apime_AstCCC -----------SYWKQCAFNAVSCFG
Nasvi_AstCCC -----------NYWRQCAFNAVSCFG
Pedhu_AstCCC -----------SYWKQCAFNAVSCFG
Dappu_AstCCC -----------SYWKQCAFNAVSCFG
Rhimi_AstCCC -----------SGWKQCSFNAVSCFG
```

Bursicon alpha and bursicon beta

```
Drome_bursalpha -QPDSSVAATDNDITHLGDDCQ-VTPVIHVLQY---------PGCVPKPIPSFACVGRCASYIQVSGSKIWQMERSCMC
Bommo_bursalpha ----------HEVQLPPGQECQ-MTAVIHVLKH---------RGCKPKAIPSFACIGKCTSYVQVSGSKIWQMERTCNC
Trica_bursalpha LDPRLNS-KIQVSGASTTDECQ-VTPVIHVLQY---------PGCVPKPIPSFACIGRCASYIQVSGSKIWQMERSCMC
Dappu_bursalpha ------------------DECQ-LTPVIHVLQY---------PGCIPKPIPSFACTGKCTSYVQVSGSKLWQTERSCMC
Rhimi_bursalpha -------------AIGPEESCQ-LRPVIHVLKQ---------PGCQPKPIPSFACHGSCSSYVQVSGSRYWQVERSCMC
Drome_bursbeta  --------LRYSQGTG-DENCETLKSEIHLIKEEFDELGRMQRTC-NADVIVNKCEGLCNSQVQPSVITPTGFLKECYC
Bommo_bursbeta  -----------------EENCETVASEVHVTKEEYDEMGRLLRSC-SGEVSVNKCEGMCNSQVHPSISSPTGFQKECFC
Trica_bursbeta  -----------VSEIS-EETCETLMSDINLIKEEFDELGRLQRIC-NGEVAVNKCEGSCKSQVQPSVITPTGFLKECYC
Dappu_bursbeta  ----------SKTNLM-SGTCETLPSTIHITKEEYTDGGILSRTC-EGDIGVAKCEGSCSSQVQPSVVHPSGFLKECMC
Rhimi_bursbeta  --TWVSASSLDNTGGG-VASCRLQETSIRITRDHSDDQGSPVRTC-EGTVLVSRCEGTCVSQVQPSITLPHGFLKECNC

Drome_bursalpha CQESGEREAAVSLF-CPKVKPGER-----KFKKVLTKAPLECMCRPCTSIEESGIIPQEIAGYSDEGPLNNHFRRIALQ
Bommo_bursalpha CQESGEREATVVLF-CPDAQNEEK-----RFRKVSTKAPLQCMCRPCGSIEESSIIPQEVAGYSEEGPLYNHFRKSL--
Trica_bursalpha CQESGEREASVSLF-CPKAKPGER-----KFIKVTTKAPLECMCRPCTGVEESAVIPQEIAGYADEGPLSNHFLKSHSQ
Dappu_bursalpha CQESGEREATVSLL-CPKAAPGEP-----KLRRVVTRAPVDCMCRPCTALEESAVMPQEIARFLDDGSFPFKL------
Rhimi_bursalpha CQEMGEREATKAVF-CPKGP--GP-----KFRKLVTRAPVECMCRPCTAPDEASVLPQEFVGL----------------
Drome_bursbeta  CRESFLKEKVITLTHCYDPDGTRLTSPEMGSMDIRLREPTECKCFKCGDFTR---------------------------
Bommo_bursbeta  CREKFLRERLVTLTHCYDPDGIRFEDEENALMEVRLREPDECECYKCGDFSR---------------------------
Trica_bursbeta  CRESFLRERTITLTHCYDPDGVRLTAETVNSMDVKLREPAECKCYKCGDFSR---------------------------
Dappu_bursbeta  CRESFLRERVVTLTHCYDANGNRLTGK-SSSLDVKMREPADCKCFRCGDSAE---------------------------
Rhimi_bursbeta  CRETYMNRREIQLQDCFDPNGQKLMEP-DGSMTIFLEEPQDCSCHKCGG------------------------------

```

CCHa-1 and CCHa-2

```
Drome_CCHa1 --SCLEYGHSCWGAHG
Bommo_CCHa1 --SCLSYGHSCWGAHG
Trica_CCHa1 --SCLSYGHACWGAHG
Rhopr_CCHa1 --SCLSYGHSCWGAHG
Drome_CCHa2 --GCQAYGHVCYGGHG
Bommo_CCHa2 --GCSAFGHSCFGGHG
Rhopr_CCHa2 --GCSAFGHSCFGGHG
Dappu_CCHa  --NCNKYGNACFGAHG
Rhimi_CCHa  NNSCKLYGHSCLGGHG
```

Tachykinin (TK) and Natalisin (NTL)

```
Drome_TK1  -------APTSSFIGMRG
Drome_TK2  --------APLAFVGLRG
Drome_TK3  --------APTGFTGMRG
Bommo_TK1  --------IPQGFLGMRG
Bommo_TK2  --------APLGFTGVRG
Bommo_TK6  --------GQMGFFGMRG
Trica_TK1  --------APSGFTGVRG
Trica_TK5  ------MPRQAGFFGMRG
Trica_TK6  ----YPYQFRGKFVGVRG
Rhopr_TK1  ---------AMGFVGMRG
Rhopr_TK2  ------APSTMGFQGVRG
Rhopr_TK4  -------TPAMGFMGMRG
Rhopr_TK7  --------GPSGFMGVRG
Dappu_TK1  ------TPNSRAFLGMRG
Rhimi_TK2  ---------GSGFFGMRG
Drome_NTL3 DKVKDLFKYDDLFYPHRG
Drome_NTL4 --HRNLFQVDDPFFATRG
Bommo_NTL1 ------IHNEPPFWAIRG
Bommo_NTL5 -------TEENPFWANRG
Bommo_NTL8 -----SSAEDDPFYISRG
Trica_NTL1 ---ASGQEEFGPFWANRG
Trica_NTL2 --DDNDINDNEPFYVTRG
Rhopr_NTL2 GDSSSTEEVQPPFWAHRG
Rhopr_NTL3 -----DTMEQDPFWVSRG
Dappu_NTL3 --YAADGGDGVPFWATRG
Rhimi_NTL2 --HDVILEQPPGFVGARG
```

**Figure S5**: A reference guide based on multiple sequence alignments to distinguish closely-related neuropeptides. Unique amino acids within a given neuropeptide have been highlighted in color. Residues conserved across all the sequences within an alignment are highlighted in black.
